# High performance imputation of structural and single nucleotide variants using low-coverage whole genome sequencing

**DOI:** 10.1101/2023.03.05.531147

**Authors:** Manu Kumar Gundappa, Diego Robledo, Alastair Hamilton, Ross D. Houston, James G. D. Prendergast, Daniel J. Macqueen

## Abstract

**Background:** Whole genome sequencing (WGS), despite its advantages, is yet to replace standard? methods for genotyping single nucleotide variants (SNVs) such as SNP arrays and targeted genotyping assays. Structural variants (SVs) have larger effects on traits than SNVs, but are more challenging to accurately genotype. Using low-coverage WGS with genotype imputation offers a cost-effective strategy to achieve genome-wide variant coverage, but is yet to be tested for SVs.

**Methods:** Here, we investigate combined SNV and SV imputation with low-coverage WGS data in Atlantic salmon (*Salmo salar*). As the reference panel, we used genotypes for high-confidence SVs and SNVs for n=365 wild individuals sampled from diverse populations. We also generated 15x WGS data (n=20 samples) for a commercial population external to the reference panel, and called SVs and SNVs with gold-standard approaches. An imputation method selected for its established performance using low-coverage sequencing data (GLIMPSE) was tested at WGS depths of 1x, 2x, 3x, and 4x for samples within and external to the reference panel.

**Results:** SNVs were imputed with high accuracy and recall across all WGS depths, including for samples out-with the reference panel. For SVs, we compared imputation based purely on linkage disequilibrium (LD) with SNVs, to that supplemented with SV genotype likelihoods (GLs) from low-coverage WGS. Including SV GLs increased imputation accuracy, but as a trade-off with recall, requiring 3-4x depth for best performance. Combining strategies allowed us to capture 84% of the reference panel deletions with 87% accuracy at 1x depth. We also show that SV length affects imputation performance, with provision of SV GLs greatly enhancing accuracy for the longest SVs in the dataset.

**Conclusions:** This study highlights the promise of reference panel imputation using low-coverage WGS, including novel opportunities to enhance the resolution of genome-wide association studies by capturing SVs.

## Background

The use of whole genome sequencing (WGS) is increasing due to reducing costs and accessibility of sequencing technologies. Nonetheless, WGS, performed at the depths required to accurately detect genetic variants, remains more expensive than strategies that genotype a reduced set of variants, typically single nucleotide variants (SNV), including arrays [1] and reduced representation sequencing [2]. Consequently, WGS remains less commonly used for large-scale genotyping applications such as genome-wide association studies (GWAS). WGS has major advantages though, in theory capturing all SNVs and indels in each sample, enhancing the chance of identifying causative variants for a target phenotype, rather than relying on linkage disequilibrium (LD) of nearby markers [3]. WGS also enhances scope to directly identify and genotype other classes of variation, such as larger structural variants (SVs) [4], although the accuracy of such calls is generally substantially lower than for SNVs when using the most common short-read sequencing approaches.

Several frameworks exist for statistical imputation of genotypes in individuals with limited genotyping information, exploiting data from individuals that have been densely genotyped [5–7]. Imputation can improve statistical power in GWAS and increase resolution to fine map causative variants in quantitative trait loci (QTL) regions [5–7]. Using closely related individuals with known pedigrees, densely typed genotypes for a few individuals (e.g. parents) can be accurately imputed to related individuals (e.g. offspring) based on shared haplotypes [5]. It is also possible to apply imputation using broader reference panels of genetic variation that capture different populations and haplotypes within a species, as done routinely in humans (e.g. [8]). Within this framework, methods have been developed for imputation against a reference panel using low coverage WGS data [9–13]. Imputation from low coverage short-read sequencing data has the advantage of reduced sequencing costs, while sufficient alleles are still covered to effectively reconstruct haplotypes in the individual. In recent studies using the most advanced algorithms in humans, this strategy was shown to have the potential to be more accurate and cost-effective at defining sample genotypes than array-based imputation against a reference panel [12,13].

To date, imputation against a species-level reference panel has focussed mostly on SNVs and indels, largely ignoring SVs, except for limited work in humans [7,14]. While SNVs and indels are markedly easier to detect and genotype with high accuracy, SVs are thought to contribute more to phenotypic variation, including by influencing gene expression with larger effect sizes [15–18] and showing higher enrichment in GWAS haplotypes [19]. Therefore, imputation of SVs using low coverage WGS should markedly increase the ability of GWAS to identify causative variants.

Atlantic salmon (*Salmo salar*) is a species of great societal importance, being among the top farmed aquaculture species globally by value. Furthermore, wild populations have major conservation value and cultural significance [20,21]. To support genetic studies with both farmed and wild Atlantic salmon populations, numerous genomic resources have been developed for this species [20,22]. This includes reference genomes for different phylogenetic lineages [23–25], high-density SNV arrays (e.g. [26,27]), and WGS data spanning diverse populations (e.g. [24,28,29]). Atlantic salmon benefits from an existing panel of high-confidence SVs detected using WGS across 492 individuals representing European and North American populations [29] and a global set of SNVs from most of the same individuals [30]. These past efforts provide a broad representation of both SNVs and SVs to leverage as a reference panel for imputation studies.

Here, the primary aim was to investigate the potential for joint SV and SNV imputation using low-coverage WGS imputed against haplotypes phased from an existing, high-quality reference panel. Data and resources available on Atlantic salmon were used as a test-case but results should apply to other species with similar resources. To achieve this aim, we utilized GLIMPSE [12] - an imputation method that, to date, has been used to impute SNVs using low-coverage WGS. Our results describe the performance of GLIMPSE for imputation of both SNVs and SVs at four WGS depths. We discuss the potential of such approaches to enhance the resolution of downstream GWAS and better characterise the role of SVs in shaping economically and ecologically important phenotypes.

## Methods

### Reference panel of SNVs and SVs

Our reference panel integrated SV and SNV genotypes from the same wild Atlantic salmon individuals (n=365) [see Additional file 1, Table S1]; mean WGS depth=8x, min depth=4.2x, max depth= 17.5x) from two independent studies using the ICSASG_v2 genome [23]. These individuals represent diverse European populations from a broad geographical range [29]. For SVs, this included 13,999 deletions, 1221 duplications, and 241 inversions that were curated individually using a method shown to exclude the majority of false positive calls while retaining mainly true variants [29]. The reference panel included 3,156,297 high quality SNVs [30]. SVs and SNVs were merged into a single VCF file using Picard Toolkit v2.23.8 [31] and then sorted and indexed using BCFtools v1.12 [32].

### Overview of imputation strategy

The key aim of this study was to test the ability of GLIMPSE v 1.1.1 [12] to impute SNVs and SVs in Atlantic salmon. This section outlines the general workflow and its rationale, followed by detailed methods in sections that follow. Code for the full workflow is available through GitHub (https://github.com/manugundappa/SV_imputation_pipeline).

GLIMPSE was tested against the two sets of samples that are described in Additional file 1, Table S2 and Additional file 1, Table S3. The first set represents samples taken from within the reference panel (hereafter: ‘in-panel’), comprising n=98 individuals, of which half were from various rivers in North Norway and the remaining half from various rivers in South Norway, with an overall mean and standard deviation (s.d.) for WGS depth of 7.8x and 1.49x, respectively [see Additional file 1, Table S2]. These samples were dropped from the training panel in-turn when assessing imputation performance. The second set represents samples external to the reference panel (hereafter: ‘out-panel’), comprising 20 farmed individuals from the Hendrix Genetics Landcatch Strain (Norwegian origin), with mean and s.d. WGS depth of 15.7x and 1.1x, respectively [see Additional file 1, Table S3]. This included two individuals from each of ten families that were previously shown to have contrasting within-family genomic estimated breeding values for resistance to amoebic gill disease [33]. The out-panel WGS data was generated as described in a following section.

Using the in-panel samples provided a benchmark to test the performance of SNV and SV imputation, where the variants were known to match with haplotypes that were phased in the reference panel. The out-panel samples were used to test imputation performance for samples that were not represented within the reference panel, which consists largely of wild individuals sampled from locations across Norway, inclusive of rivers from which the founders of Norwegian aquaculture were derived [34]. The Hendrix-Genetics Landcatch strain we used was derived from sources with strong connections to the founder populations of Norwegian aquaculture and, hence, was expected to also share haplotypes with the reference panel. A further rationale for choosing samples from a commercial aquaculture population was to demonstrate utility of the developed pipeline in commercial aquaculture breeding programs.

The GLIMPSE workflow was validated for and by default restricted to SNVs [12]. We modified it to accommodate SVs and provide an efficient workflow to jointly impute SNVs and SVs (Fig. 1). Briefly, both the in-panel and out-panel datasets were down-sampled to 1x, 2x, 3x, and 4x depths. The reference panel was processed to generate a list of all variable SNV and SV positions for which imputation should be performed. Genotype likelihoods (GLs) for SNVs within the reference panel were generated for each down-sampled WGS dataset as input for the GLIMPSE pipeline (see below). Imputation was firstly performed against the down-sampled datasets across all variable SNVs and SVs by only using the SNV GLs. We also developed and tested an alternative approach in which the pipeline was provided SV GLs (see below). The imputed SNV and SV genotypes were later compared to reference genotypes derived from the original higher coverage WGS depth data to calculate imputation performance.

**Figure 1.**
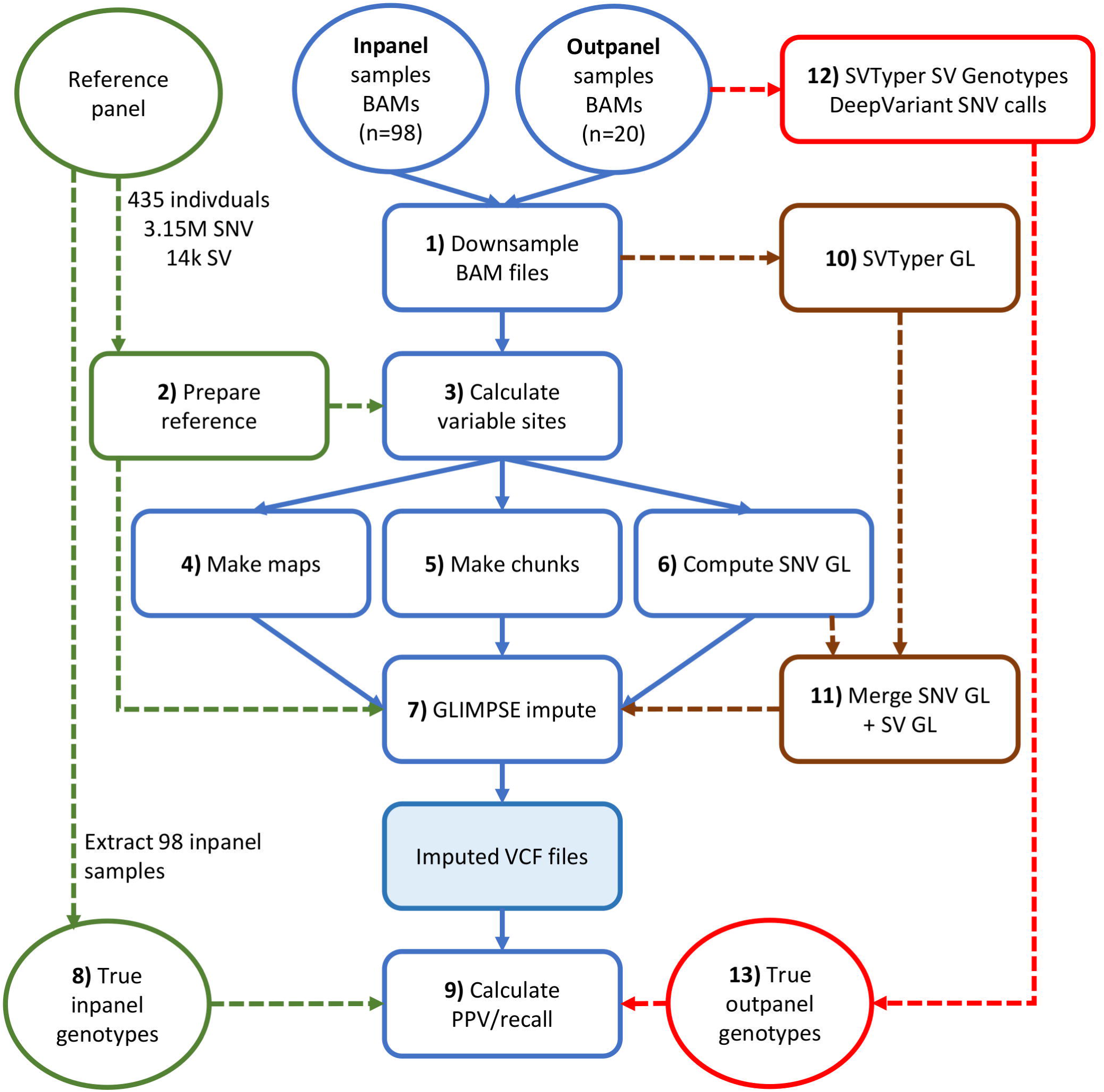
Work flow describing steps of the imputation pipeline. Highlighted in blue is the core GLIMPSE pipeline we used to impute SNVs and SVs. Steps in green highlight the details of the reference panel used for imputation, in brown the additional steps used when running GLIMPSE supplemented with SV genotype likelihoods, and in red are steps highlighting generation of SNV and SV genotypes for the out-panel samples and downstream use in the imputation pipeline. Additionally, during imputation of in-panel samples (n=98), the reference panel was updated using a ‘leave one-out’ strategy, removing one individual to be imputed in each round of imputation (i.e. process repeated 98 times). PPV = Positive predictive value, GL = Genotype Likelihood.

### Imputation of in-panel samples

#### GLIMPSE imputation ignoring SV GLs

Samtools v1.12 [32] *view* was used to down-sample BAM files per chromosome. The Samtools *coverage* command was used to compute the mean sequencing depth and down-sampling factors. The Samtools *view* command was used to down-sample the BAM file for each chromosome to the target depths (1x, 2x, 3x, 4x) using the computed down-sampling factors. The down-sampled BAM files were used as inputs to GLIMPSE (Fig. 1: Step 1).

BCFtools was used to split the reference panel by chromosome and to exclude one of the 98 individuals from the reference panel. This was done in turn such that each sample was excluded individually, with the other 97 samples remaining in the training panel (Fig. 1: Step 2). The output of this step was stored in a BCF file. Variable positions across each chromosome-specific BCF file were identified using the BCFtools *view* command (Fig. 1: Step 3). Although the GLIMPSE pipeline restricts this step to SNVs, considering our objective of imputing SVs, we ignored the *–v snps* flag during this step. A proxy genetic map was generated across all samples and chromosomes (assuming 1Mb per cM) using the variable sites information and a custom R script (Fig. 1: Step 4). The GLIMPSE_*chunk* command was then used to define regions for which to perform imputation and phasing across all chromosomes. A window size of 100 Mb and buffer size of 200 kb was used for this step (Fig. 1: Step 5). GLs were calculated for all variable SNVs within the variable sites datasets using the *mpileup* command within BCFtools (Fig. 1: Step 6). SV GLs were not included in this step, as the objective was to impute SVs using GLs for SNVs only (i.e. on the basis of LD). Finally, all variants, including SNVs and SVs, were imputed against the reference panel using the GLIMPSE_*phase* command at the four down-sampled WGS depths (Fig. 1: Step 7). This command generates a VCF file containing genotype probabilities based on a dataset that contains the low depth GLs, the reference panel of all haplotypes, the genetic map, and chunk information across each sample and chromosome to. BCFtools *view* was used to extract the original reference genotypes of all variants from the reference panel that were previously called from the higher coverage sequence data using GATK [30] for comparison against the imputed genotypes (Fig. 1: Step 8).

#### GLIMPSE imputation accuracy estimation

The imputed genotype values for variants were extracted and filtered using a custom R script to calculate imputation accuracies in comparison to the reference genotypes (Fig. 1: Step 9). Imputed VCF files for each sample were merged across all chromosomes and processed to extract posterior support values for each variant. We then filtered the imputed variants (converting imputed genotypes to missing values ‘./.’) at different genotype posterior probability cut-off values of 0.60, 0.75, and 0.90, in order to establish the trade-off between imputation accuracy and recall rate (defined below). We then compared the imputed genotypes against the higher-coverage genotype calls in the reference panel to generate a matrix of ‘true’ vs imputed genotypes for SNVs and different categories of SVs. The output for each sample at different depths was used to generate positive predictive values (PPVs) and recall rates, to gauge overall imputation performance. PPV was computed as the ratio of the number of correct genotype calls divided by the sum of correct and incorrect genotype calls (PPV = number of “correct” SV genotype calls / (number of “correct” SV genotype calls + number of “incorrect” SV genotype calls)). Recall rate was calculated as the proportion of variants imputed against the total number of variants in the reference panel (Recall rate = number of SVs imputed / total number of SVs). PPVs and recall rates across all samples and posterior cut-offs were visualised using the package ggplot2 [35] within R.

#### GLIMPSE imputation including SV GLs

We devised an approach to feed SV GLs into the GLIMPSE imputation pipeline and test its impact on SV imputation performance. SVTyper [36] was run via the Smoove [37] pipeline to genotype all SVs in the reference panel across all down-sampled BAM files for the ten samples (Fig. 1: Step 10). A custom R script was used to convert SV GLs into phred-scaled genotype likelihoods (PLs), rounded to the closest integer. The VCF files containing SV PLs were then merged with the SNV GL VCF files that were generated in the GLIMPSE pipeline using Picard toolkit v2.23.8 [31] (Fig. 1: Step 11). The rest of the pipeline remained the same as described when running imputation without SV GLs, except for running the GLIMPSE_phase command after removing the *--impute-reference-only-variants* flag. The final imputed genotypes were processed as described above to generate PPVs and recall rates.

#### Impact of SV length on imputation accuracy

Unlike SNVs, which uniformly impact a single base pair in the genome, SVs cover a broad spectrum of lengths, which may impact the accuracy of SV imputation. To test this, we explored imputation results from 1x and 4x depths, separately for each SV class, across the 98 in-panel samples. We restricted this test to posterior probability cut-offs ≥0.90 to ensure we retained only high-confidence imputation calls. To evaluate the PPVs for different lengths, SV lengths were split into bins of 100-500 bp, 500-1000 bp, 1000-2000 bp, 2000-5000 bp, 5000-10,000 bp, and 10,000-2,000,000 bp. To this end, our custom script to summarise imputation accuracies (described above) was modified to filter for minimum and maximum SV lengths. PPVs for GLIMPSE only and GLIMPSE including SVTyper genotype likelihoods were plotted using ggplot2 on R.

#### Merging SVs across different imputation strategies

Supplying the GLIMPSE pipeline with SV GL’s increased PPV but reduced recall rates (see Results). We therefore developed an approach to merge the outputs from both approaches (i.e. either including or excluding SV GLs during imputation) to test if performance was optimized by taking the ‘best of both’ strategies. A custom R script was used to overlap the VCF files obtained from both the pipelines. For SVs, we tested two distinct approaches for downstream processing, defined as ‘logical’ and ‘non-logical’ filtering. Logical filtering took forward SV genotypes with a higher likelihood, while non-logical filtering took forward SV genotypes from GLIMPSE imputation that included SV GLs, irrespective of whether the alternative approach (i.e. excluding SV GLs) had a better likelihood. After applying these filters, we merged the output with all SVs imputed by the approach that excludes SV GLs, making a comprehensive dataset of all imputed SVs from both approaches. We then calculated PPVs and recall rates as described above.

### Out-panel SV and SNV detection and genotyping

DNA was extracted from each sample using a standard phenol chloroform method. Purity of the DNA was confirmed using Nanodrop 1000 (Thermo Scientific) and the integrity was confirmed using agarose gel electrophoresis. Approximately 1.0μg DNA per sample was used to generate the DNA sequencing libraries by Novogene using the NEBNext® DNA Library Prep Kit. The DNA libraries were sequenced on an Illumina® Novaseq 6000 platform by Novogene UK to generate paired-end 150 bp reads (information on sequencing data provided in Additional file 1, Table S3).

Raw fastq data were quality checked using fastqc [38] and trimmed to remove adapters and low quality bases (PHRED score<25) using Trimgalore (default parameters) [39]. The trimmed reads were aligned to Atlantic salmon genome version ICSAG_v2 (NCBI accession: GCA_000233375.4) using BWA [40] to generate alignment files (SAM format), which were converted into BAM format using Samtools [32]. SV calling was performed using the pipeline described in Bertolotti et al. (2020) [29]. Briefly, the alignment BAM files were processed using the SMOOVE v2.3 pipeline, which uses Lumpy v 0.2.13 [37] to generate SV calls across all 20 samples (Fig. 1: Step 12). Gap regions and high coverage regions in the genome were excluded from SV calling [29]. A VCF file that merged all *de novo* SV calls was manually curated using SV-plaudit [41] to generate a list of high-confidence SV calls. This final set of out-panel SVs was overlapped with the reference panel SVs generated by Bertolotti et al. (2020) [29] using Bedtools *intersect*. Only SVs with a strict reciprocal overlap of 90% across both datasets were considered shared by the out-panel and the in-panel for further analyses. Histograms summarizing SVs lengths were generated using ggplot2 [35]. SV overlap with SVs from [29] was illustrated using the package VennDiagram in R [42].

SNV calling was performed using the BAM alignments. DeepVariant [43] was used to call SNVs, setting *--model_type=WGS*, with other parameters as default (Step 12). The output gVCF files were split by chromosome using the BCFtools *view* command. GLnexus [44] was used (default parameters) to perform joint variant calling and to merge gVCFs from all samples by chromosome. This was done with the config set to ‘DeepVariantWGS’ to produce an individual merged BCF file for each chromosome. BCF files from all chromosomes were merged using the BCFtools *concat* command. Vcftools [45] was used to filter SNVs using the following additional criteria: *--min-alleles 2 --max-alleles 2 --maf 0.01 --max-missing 0.7 -- remove-indels --minGQ 10 --minDP 4 --recode --recode-INFO-all --minQ 30.* SNVs from the out-panel samples were merged with those from the reference panel to generate a PCA plot and a genomic relationship matrix. The merged VCF file was processed using PLINK [46] to calculate eigenvalues of the genetic covariance matrix across all samples and a PCA plot of PC1 versus PC2 was generated using ggplot2. Genomic relationships for all samples in the merged VCF file were also generated using PLINK, based on the entire SNV dataset. The mean relationships of the 20 out-panel samples against the in-panel samples from North Norway, South Norway, and the entire reference panel were plotted in R using the ggplot2 package.

### Imputation of out-panel samples

SV and SNV imputation was performed for the 20 out-panel samples as described above (i.e. Fig. 1), with the following modifications: the reference panel included all 365 individuals in Step 2; GLIMPSE phase was run across all samples to impute SNVs and SVs against the reference panel (Step 7); imputation of SVs in the out-panel samples was performed by generating SVTyper GLs for SVs in the reference panel (Step 10) and by merging them into the pipeline (Step 11); ‘true’ genotype calls were extracted from the DeepVariant SNV calls and SVTyper genotypes derived from the 15x data (Steps 12-13). PPVs and recall rates were calculated and visualized as described for the in-panel samples.

## Results

### In-panel imputation performance for SNVs

We first assessed the impact of sequencing depth and posterior probability (PP) cut-off on SNV imputation performance. Increasing sequencing depth is expected to increase the accuracy of downstream genotype calls, but at increased financial cost. Likewise, increasing the PP cut-off is expected to lead to more accurate calls among the remaining genotypes, but at the trade-off of having less genotype calls remaining that pass filtering. SNVs were found to be imputed with high accuracy for the in-panel samples, irrespective of sequencing depth or PP cut-off. PPVs ranged from PPV = 0.96 at 1x depth with PP > 0.6, to PPV = 0.99 at 4x depth with PP > 0.90 (Fig. 2a). As expected, higher PP cut-offs increased SNV imputation accuracy across all WGS depths, but at the expense of lower recall rates (Fig. 2a, b). Recall rates varied markedly by WGS depth and cut-off (Fig. 2b). The highest recall rate (81%) was achieved at 4x depth with PP > 0.60 (PPV = 0.98). Recall rate increased slightly with WGS depth when using PP > 0.60 (from 0.77 to 0.81 at 1x to 4x depth). Using PP > 0.90 resulted in recall rates from 61 to 75% at 1x to 4x depths. We also tested whether there was any difference in PPV and recall rates across the samples from the North and South of Norway, which are known to be genetically distinct [29]. PPV was the same for these two sets of samples across all PP value cut-offs [see Additional file 2, Fig. S1-S2]. However, there was a marginal increase in recall for North compared to South Norwegian samples [see Additional file 2, Fig. S1 and Additional file 2, Fig. S2]. In summary, GLIMPSE showed excellent performance for SNV imputation using the in-panel samples, capturing 1,920,956 SNVs [see Additional file 1, Table S4] with 98% accuracy (i.e. PPV = 0.98, recall rate = 0.62) at 1x depth when applying the PP=0.9 cut-off.

**Figure 2.**
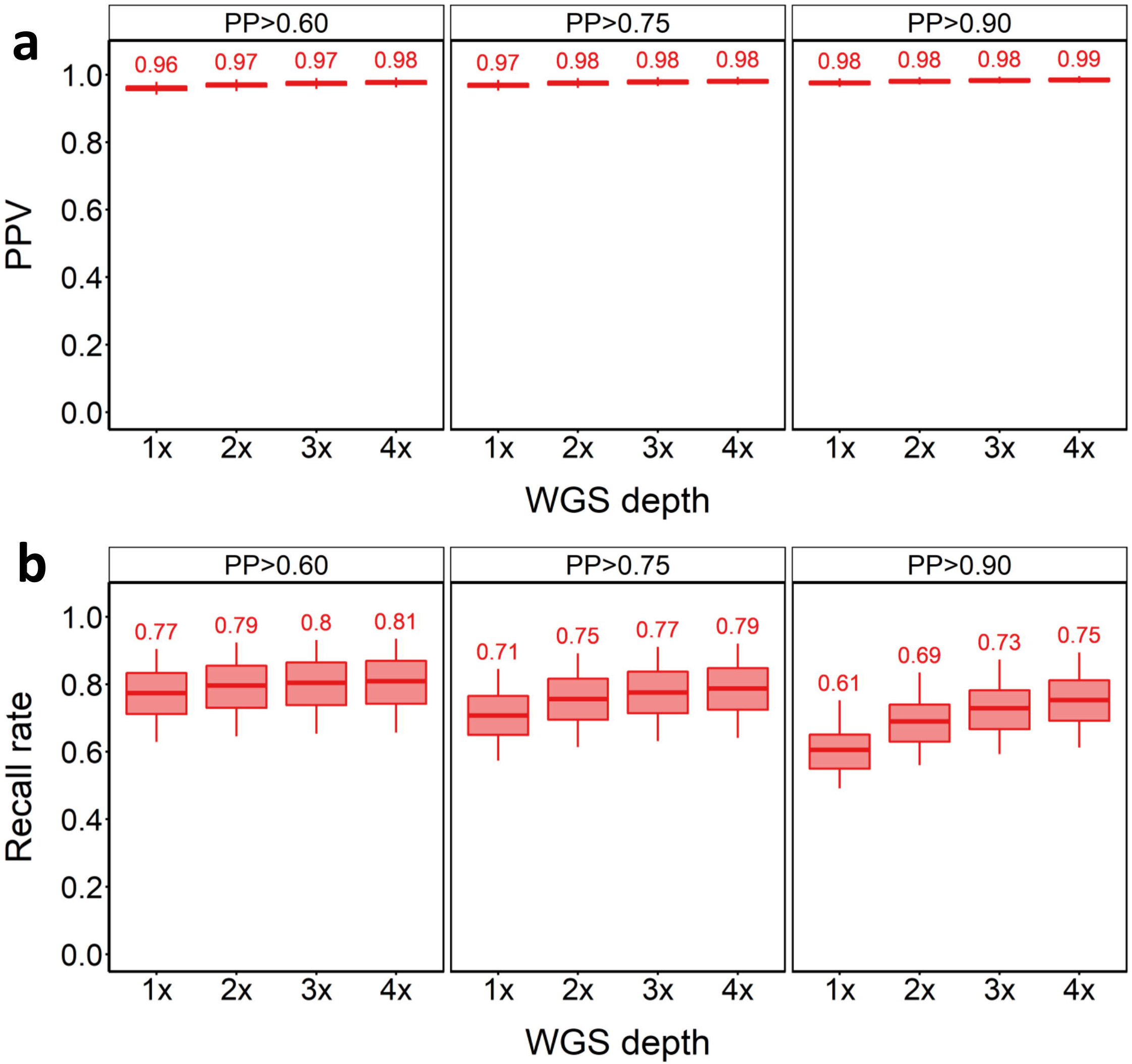
SNV imputation results for in-panel samples. PPV (**a**) and recall rate (**b**) for SNVs across tested WGS depths and PP cut-offs.

### In-panel imputation performance for deletion SVs

We next assessed the ability to impute genotypes at deletion SVs using GLIMPSE. We compared two approaches. In the first, we ignored reads in the sequencing data that overlap an SV, relying solely on LD between SNVs and SVs to impute the deletion genotypes. In the second approach we provided GLIMPSE with prior SV GLs for deletions that were called using SVTyper (‘GLIMPSE+SVtyper’ approach), therefore accounting for reads observed at SV locations in the corresponding low-depth WGS data. As evident from Fig. 3a, the GLIMPSE+SVtyper approach consistently increased deletion imputation accuracy compared to relying on LD between SNVs and SVs alone. This demonstrates that accounting for the low coverage sequencing reads at SV positions improves the SV calls. However, applying this strategy introduces a trade-off of lower recall rates, which was particularly evident with very low coverage WGS (Fig. 3b). As GLIMPSE will only make calls for SVs with prior GLs when run using the GLIMPSE+SVTyper approach, any SVs that lack prior GLs (i.e. owing to lack of sequencing data) are excluded. In contrast, when run without prior SV GLs, GLIMPSE will call genotypes at all SVs, increasing the corresponding recall rates. At 4x depth, a maximum recall rate of 90.3% (PP > 0.6) was achieved for the GLIMPSE+SVTyper method, approaching the recall rate using GLIMPSE without supplementation of SV GLs (97.2%) (Fig. 3b), although with a higher PPV than when relying on LD alone (0.84 vs 0.75). While samples from the South of Norway achieved 90.1% recall using GLIMPSE+SVTyper at 4x depth, GLIMPSE-only achieved 97% recall [see Additional file 2, Fig. S3]. In-panel samples from the North of Norway performed marginally better than those from South Norwegian samples, achieving, respectively, 90.5 and 97.2% recall (GLIMPSE only vs GLIMPSE+SVTyper) at 4x depth [see Additional file 2, Fig. S4]. However, PPVs remained nearly same compared to those obtained from the merged in-panel outcome for both the Northern and Southern Norwegian samples (see Additional file 2, Fig. S3-S4).

**Figure 3.**
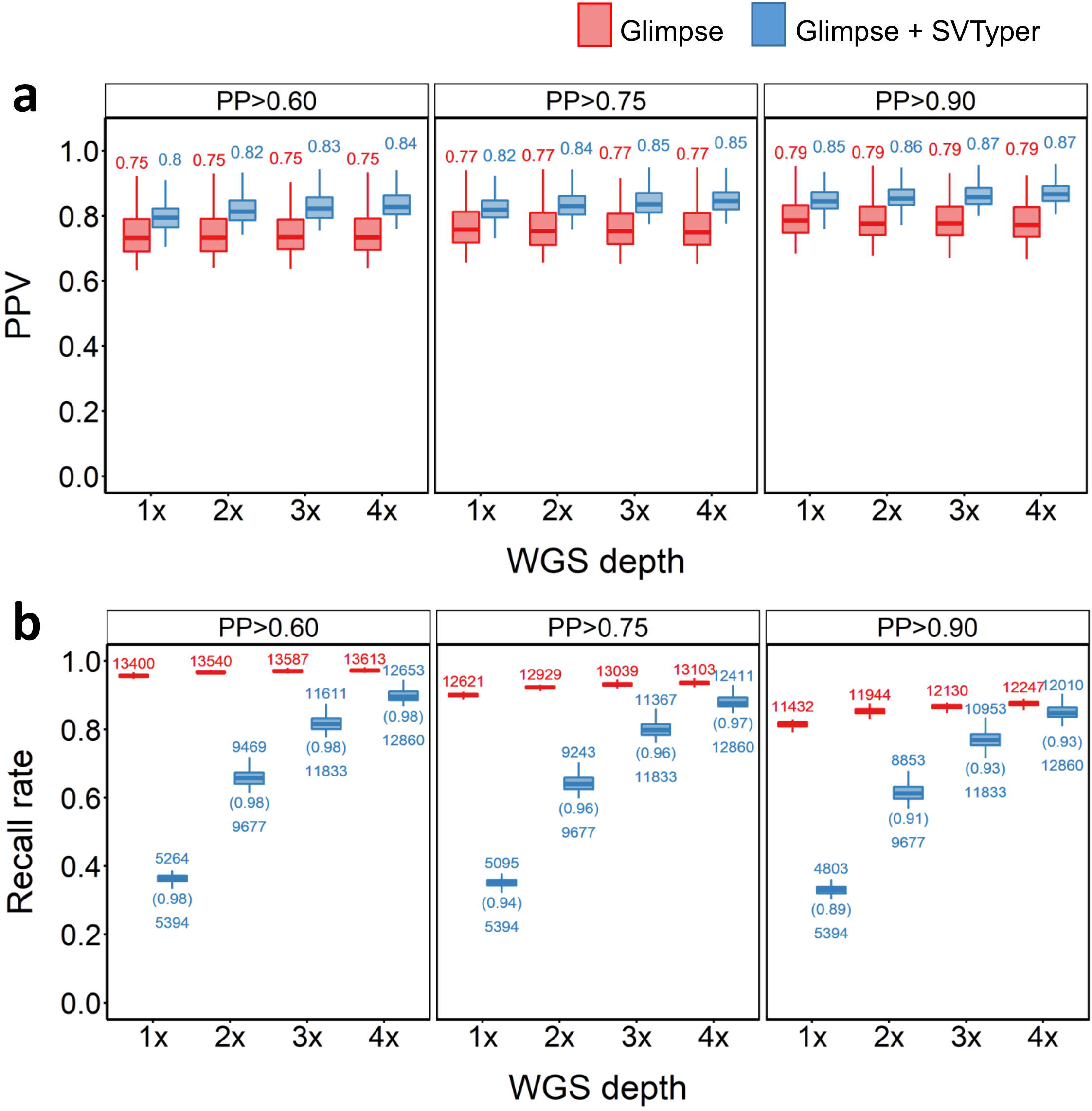
SV deletion imputation results for in-panel samples. a) PPV (**a**) and recall rate (**b**) for across different PP cut-offs, sequencing depths and strategies used. Red and blue plots indicate PPVs when the pipeline was supplied with SNP GLs only, or both SNP and SV GLs, respectively. The numbers above the box plot indicate the number of SVs that were imputed, the numbers below the box plots indicate the number of SVs with genotype likelihood that were supplied into the GLIMPSE pipeline and the numbers in parenthesis indicate the recall rate based on the number of SVs fed into the pipeline.

### In-panel imputation performance for duplication and inversion SVs

PPVs were lower for duplication and inversion SVs than for deletions, irrespective of whether GLs were supplied to GLIMPSE [see Additional file 2, Fig. S5-S6]. For both SV types, we observed higher PPVs using GLIMPSE+SVtyper, with a stronger effect than for deletions [see Additional file 2, Fig. S5-S6]. As for deletions, using GLIMPSE-only, WGS depth had little impact on PPVs and recall across the different WGS depths and PP cut-offs for both SV classes [see Additional file 2, Fig. S5-S6]. Compared to deletions, there was more sample-to-sample variation in PPV and recall for duplications and inversions across different WGS depths and PP cut-offs [see Additional file 2 Fig. S5-S6]. As for deletions, the highest PP cut-off (PP=0.9) decreased recall across both imputation strategies for duplications and inversions [see Additional file 2 Fig. S5 and Additional file 2 Fig. S6].

When using GLIMPSE+SVTyper for duplications, the highest PPVs were achieved at 1x depth across all tested PP cut-offs [see Additional file 2 Fig. S5]. This is likely explained by the extremely low recall at 1x depth [see Additional file 2 Fig. S1], suggesting a subset of SV duplications captured at 1x depth is biased compared to a set that imputes accurately. When using GLIMPSE+SVTyper, recall of both duplications and inversions increased with WGS depth, reaching maximum values of 85.3 (PP > 0.6) and 91% (PP > 0.6), respectively, at mean 4x depth [see Additional file 2, Fig. S5-S6]. The best overall performance for both SV classes was observed at 4x depth with PP > 0.9 with, respectively, 75.2 and 83.3% of the duplication and inversion SVs captured with PPVs of 0.77 and 0.82, respectively [see Additional file 2, Fig. S5 and Additional file 2, Fig. S6]. Duplication and inversion SVs analysed separately for North and South Norwegian samples revealed little differences in terms of PPVs and recall; the South samples were associated with lower recall and a slightly higher PPV value compared to North samples [see Additional file 2, Fig. S7-S10].

### Does SV length impact SV imputation?

While our results demonstrate that accurate SV imputation is possible using GLIMPSE, even at low sequencing depths, it remained unclear whether variation in SV lengths, which does not occur for SNVs, influences the accuracy of SV imputation, and if so, whether this depends on sequencing depth. We therefore investigated the accuracy of SV imputation at 1x and 4x depths for all in-panel samples, separately for each SV class but restricted to imputed SV calls with PP ≥0.90.

For both GLIMPSE-only and GLIMPSE+SVtyper, SV imputation PPVs were highest for shorter SVs and lowest for the longest tested SV length bins, for all three SV classes [see Additional file 2, Fig. S11]. The different sequencing depths did not affect imputation accuracy when using the GLIMPSE-only approach. The GLIMPSE+SVtyper approach, which provides SV genotype likelihoods to GLIMPSE, showed higher PPVs across all SV size ranges, and in most cases, moderately increased PPVs were achieved with 4x depth, with the exception of duplications >1000 bp compared to the GLIMPSE-only approach. For deletions, GLIMPSE+SVtyper showed more consistent performance across all SV lengths than GLIMPSE-only, achieving PPVs of ∼0.8 for the largest deletions (>10,000 bp) in our dataset.

Overall, these results demonstrate that SV length influences SV imputation accuracy using GLIMPSE, supporting the value of leveraging SV genotype likelihoods to maximise imputation accuracy, even with low sequencing depth. The results also show that even large SVs can be imputed with adequate accuracy for useful downstream applications.

### Merging in-panel SV imputation results

Running GLIMPSE-only resulted in comparatively lower PPVs, but higher recall, compared to GLIMPSE+SVTyper. To exploit the benefits of both strategies, we assessed the performance of merging the SV calls from both approaches using ‘logical’ ‘and non-logical’ filtering strategies, as described in the Methods. As a general rule, logical filtering led to slightly lower PPVs and slightly higher recall rates for all SV types across different WGS depths and PP cut-offs (Fig. 4). Overall, deletions were the best performing SVs. Even at 1x depth, we were able to impute 11,815 (∼84.4%) of the reference panel deletions, with PPVs approaching 80% and with relatively small gains in accuracy achieved at higher WGS depths, but significant gains in recall, e.g. with an additional ∼1400 deletions captured at 4x coverage. PPVs of duplications and inversions were similar to results before merging, with no pronounced impact on recall (Fig. 4a, b). Similarly, the samples from North and South Norway performed similar to the merged in-panel results, with no pronounced difference in PPV and recall across all types of SVs [see Additional file 2, Fig. S12-S13].

**Figure 4.**
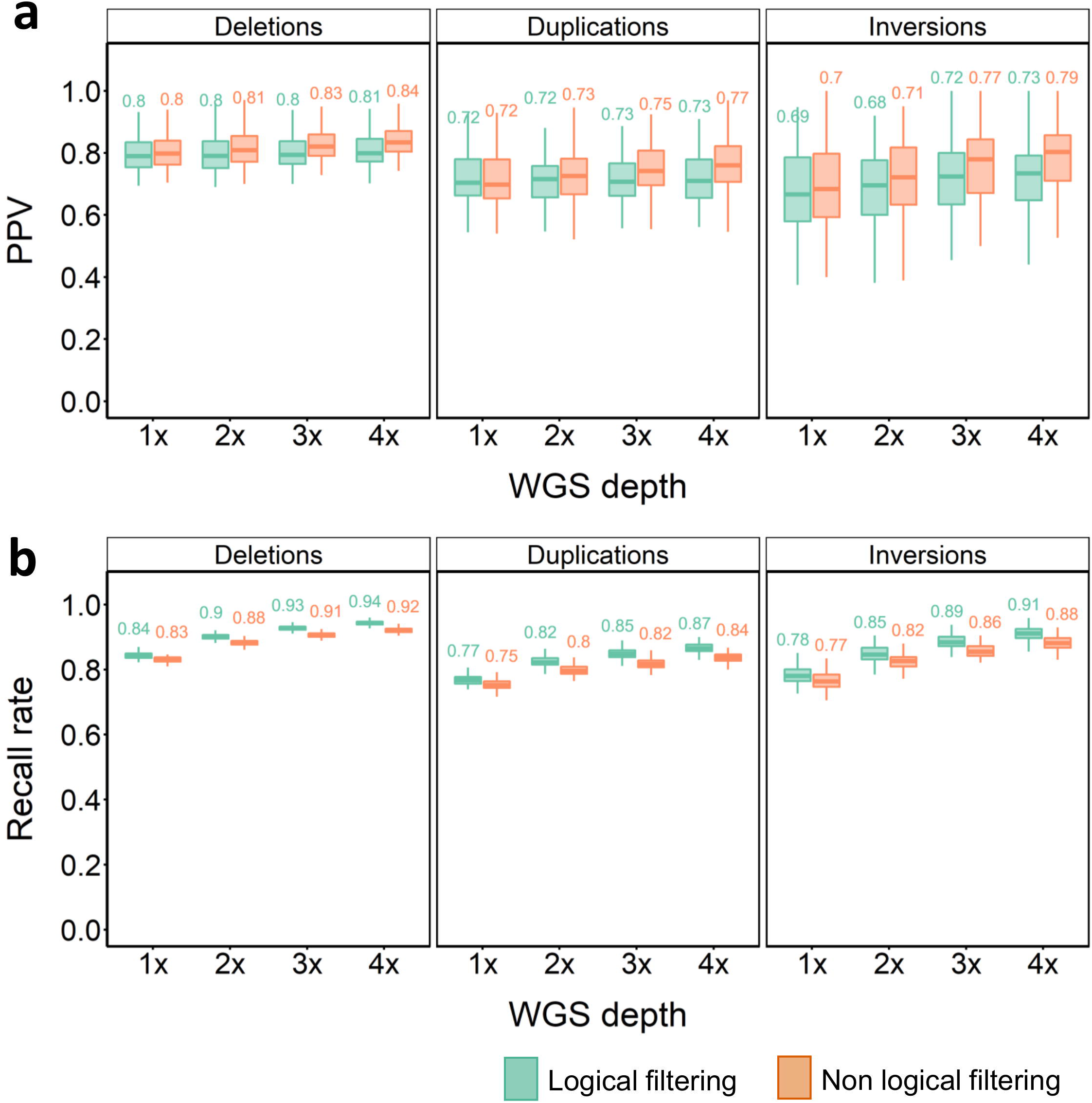
Merged SV imputation results for in-panel samples. PPV (**a**) and recall rate (**b**) for SV imputation across different SV classes, WGS depths and strategies at a PP cut-off of 0.9. Green plots indicate PPVs for the ‘logical’ and brown for ‘non-logical’ SV filtering approaches (see Methods).

### SVs and SNVs detected in the out-panel samples

To assess the performance of imputation on a set of samples that are more divergent from the reference panel, we called 44,779 SVs across 20 Atlantic salmon individuals representing the out-panel samples (35,728 deletions, 8532 duplications, 519 inversions). Manual visual curation of these variants in the SV-plaudit framework [41], using the strategy of Bertolotti et al. [29], allowed us to define a high-confidence set of 6301 SVs (6,078 deletions, 208 duplications, 15 inversions). We discarded 85.9% (i.e. 38,478/44,779) of the raw SV calls, a similar rate to that observed previously [29]. Deletions, duplications, and inversions had average lengths of 1082 bp (range: 100 – 413,436 bp, s.d. 7930 bp), 4273 bp (range: 142 – 30,608 bp, s.d. 4202 bp), and 4740 bp (range: 1152 – 16,008 bp, s.d. 4146 bp), respectively (Fig. 5a-c). Overlapping these variants against the reference panel SVs identified 3352, 96, and 3 shared deletions, duplications, and inversions, respectively (Fig. 5d-f). Thus, 2726 deletions, 112 duplications, and 12 inversions were specific to the out-panel samples (Fig. 5d-f).

**Figure 5.**
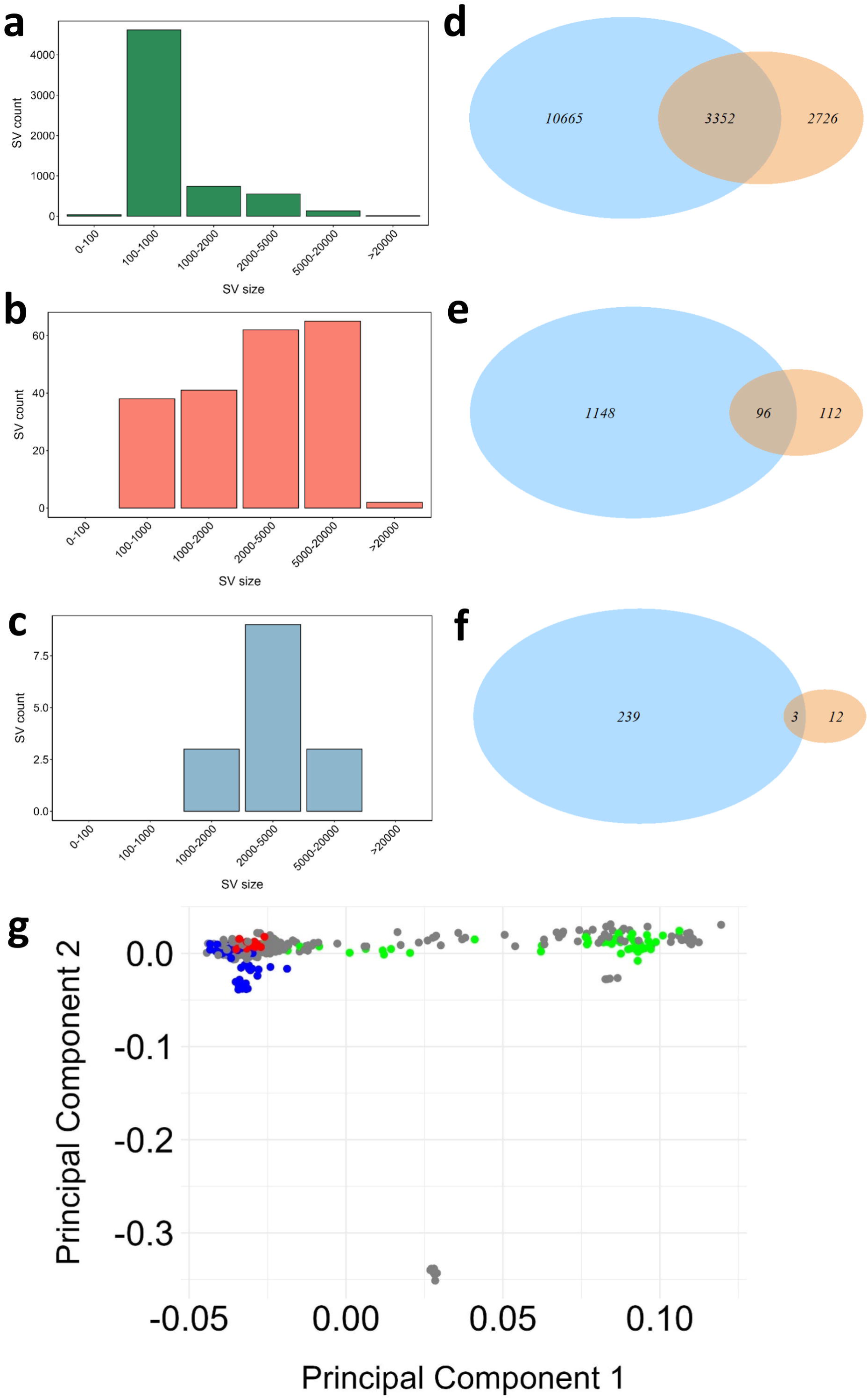
A high-confidence set of SVs detected in the out-panel samples. Size distributions are shown for deletions (**a**), duplications (**b**) and inversions (**c**). Venn diagrams highlighting overlap of SVs between the reference panel (blue circle) and out-panel sample (orange circle) are shown for deletions (**d**), duplications (**e**) and inversions (**f**). PCA plot (**g**) of all the samples in the reference panel and the out-panel samples, highlighted in red colour are the out-panel samples, blue are the in-panel South Norway samples and green are the in-panel North Norway samples.

Genotype frequencies of homozygous SVs (1/1) in out-panel samples revealed no clear difference in distribution between SVs that were unique to the out-panel population and those that overlapped the reference panel [see Additional file 2, Fig. S14]. However, counts of heterozygous SVs (0/1) were slightly higher at lower frequencies in the SVs that were private to the out-panel samples [see Additional file 2, Fig. S14].

A total of 4,009,327 SNVs were retained in the out-panel samples after applying all filters described in the methods. A PCA of the out-panel samples alongside the reference panel samples showed that the out-panel samples clustered more closely with the in-panel South Norwegian samples (Fig. 5g). The mean genomic relationships of the out-panel samples with the reference panel samples indicated no detectable recent relationship [see Additional file 2, Fig. S13] across all samples. However, the in-panel South Norwegian samples exhibited a slightly closer relationship to the out-panel samples compared to the North Norwegian samples, reinforcing observations from PCA.

### Out-panel sample imputation performance for SNVs

SNV imputation performance for the twenty out-panel samples was similar to the in-panel samples, with PPVs ranging from 0.92 to 0.98 across tested WGS depths and PP cut-offs (Fig. 6a). However, recall was markedly lower across all tested conditions (Fig. 6b), presumably owing to differences in haplotypes between the (wild) reference panel individuals and (farmed) out-panel samples. Nonetheless, we were able to impute 52 to 70% of the SNVs in the reference panel (i.e. 1.62 - 2.21 million SNVs) across the different PP cut-offs (Fig. 6b; Additional file 1, Table S4).

**Figure 6.**
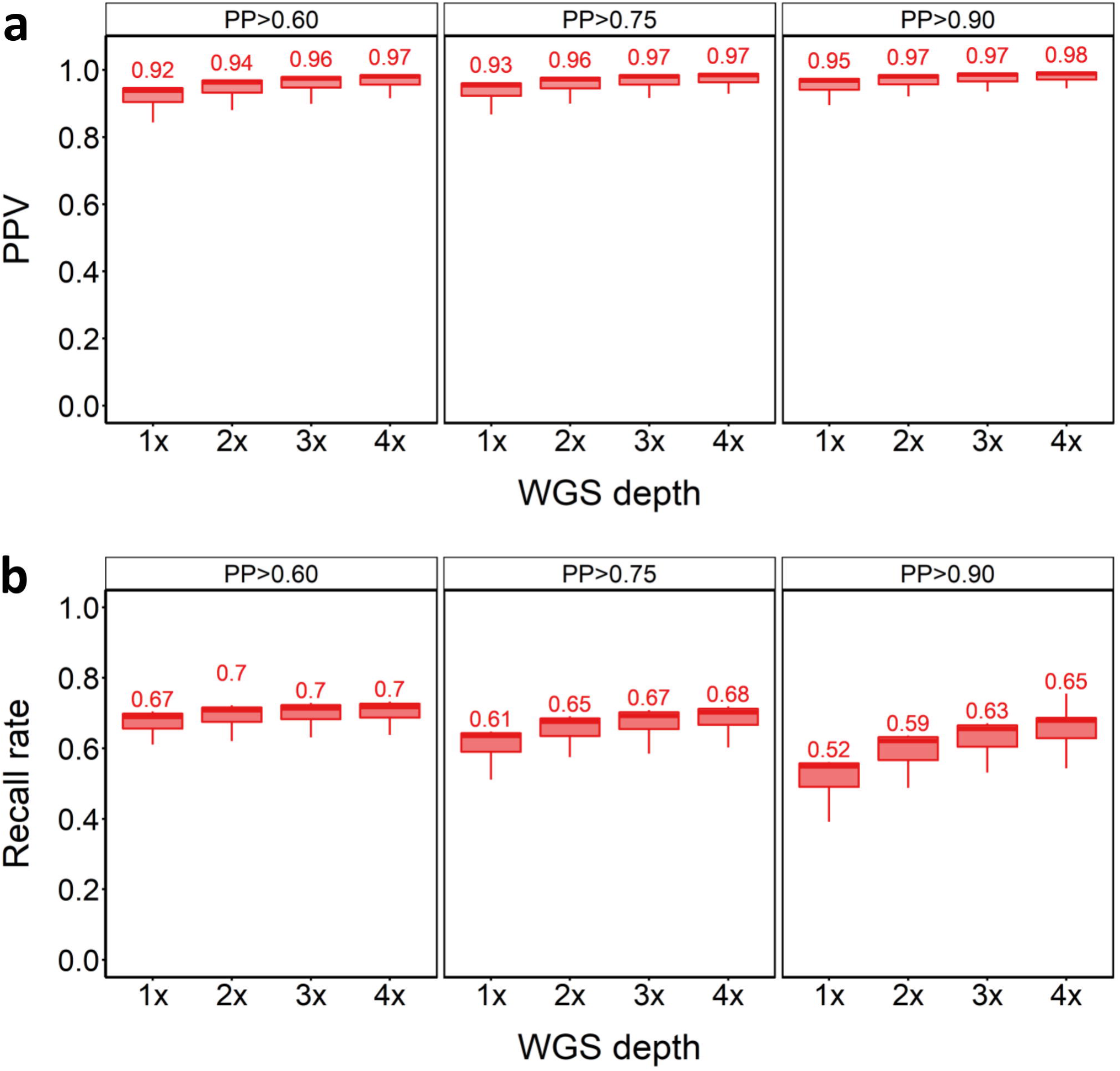
SNV imputation results for out-panel samples. PPV (**a**) and recall rate (**b**) for SNVs across tested WGS depths and PP cut-offs.

### Out-panel sample imputation performance for deletion SVs

Deletion SV imputation performance was slightly higher for the out-panel samples than for the in-panel samples (compare Fig. 3 and 7), with PPVs approaching or exceeding 0.9 using the GLIMPSE+SVtyper approach (Fig. 7a). We also observed comparatively higher PPVs for the out-panel samples using GLIMPSE-only, with PPVs >0.85 achieved even at 1x depth with PP > 0.9 (Fig. 7a). Both results are potentially driven by the greater depth of the WGS data that were used to estimate the ‘true’ genotypes in the out-panel samples. There was no major difference in recall when comparing imputation in the out-panel and in-panel samples (compare Fig. 3b and 7b). To summarise, it was possible to capture a large proportion of deletion SVs in the out-panel samples with high accuracy, even using very low WGS depths and without supplementing GLIMPSE with SV GLs.

**Figure 7.**
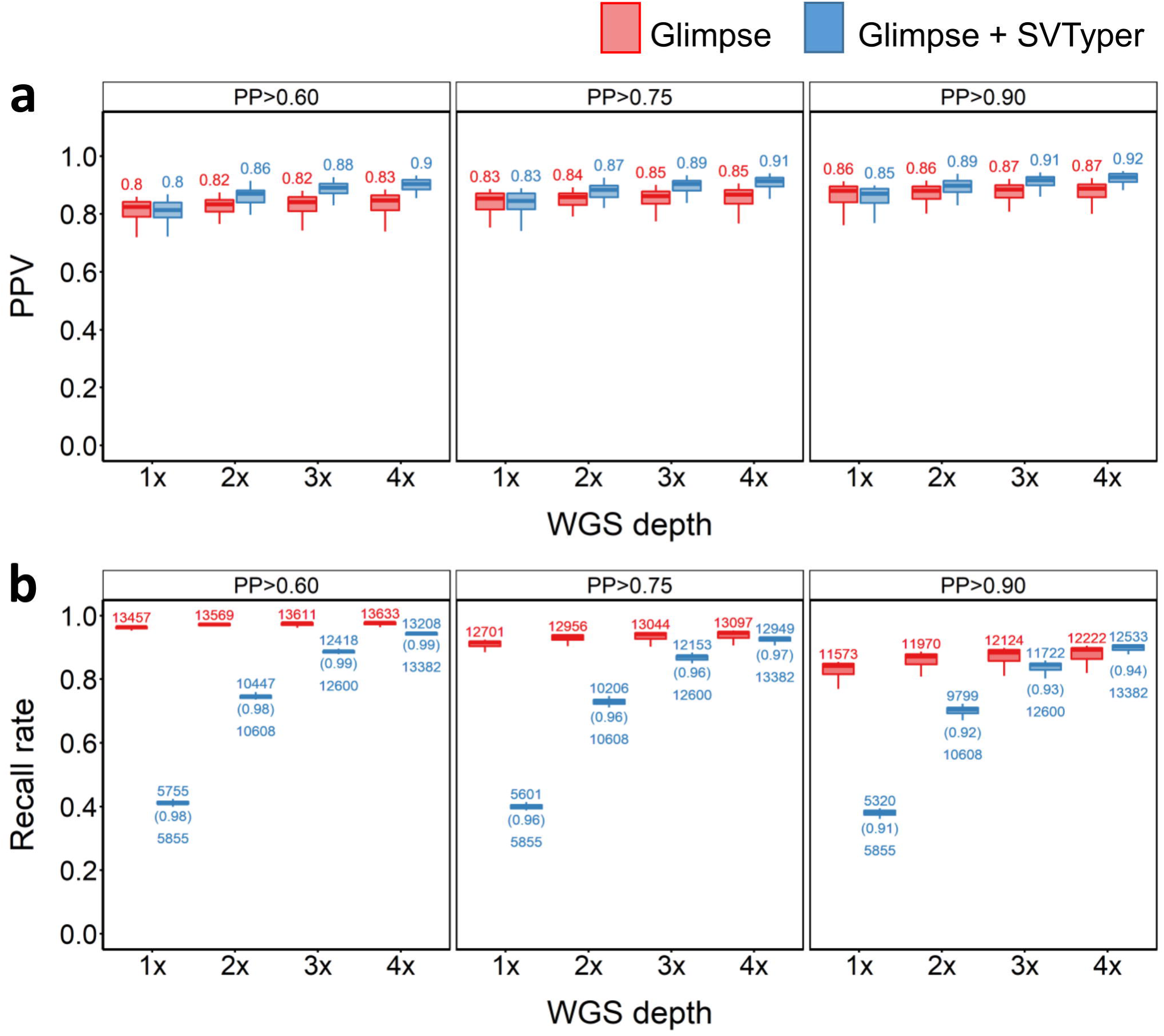
SV deletion imputation results for out-panel samples. PPV (**a**) and recall rate (**b**) for across different PP cut-offs, sequencing depths and strategies used. Red and blue plots indicate PPVs when the pipeline was supplied with SNP GLs only, or both SNP and SV GLs, respectively. The numbers above the box plot indicate the number of SVs that were imputed, the numbers below the box plots indicate the number of SVs with genotype likelihood that were supplied into the GLIMPSE pipeline and the numbers in parenthesis indicate the recall rate based on the number of SVs fed into the pipeline.

### Out-panel sample imputation performance for other SVs

PPVs for imputed duplications ranged from 0.50 to 0.60 across all WGS depths and PP cut-offs for the out-panel samples [see Additional file 2, Fig. S15], similar to the in-panel samples [see Additional file 2 Fig. S5]. As for the in-panel samples, imputation of duplication SVs performed better using GLIMPSE+SVtyper than the out-panel samples [see Additional file 2, Fig. S15]. Recall was also comparable for the out-panel and in-panel imputed duplication SVs [see Additional file 2, Fig. S15].

Only three inversions were common to the out-panel samples and the in-panel samples, although these were generally successfully imputed across coverages [see Additional file 2, Fig. S16]. With this caveat noted, inversion imputation was more accurate than observed for the larger inversion set available for in-panel samples. We achieved PPVs for inversions ranging from 0.80 to 0.90 across all WGS depths [see Additional file 2 Fig. S16a]. Recall for inversions in the out-panel samples was similar to that achieved for in-panel samples [see Additional file 2, Fig. S16b].

### Merging of out-panel sample SV imputation results

Merging outputs from the two imputation strategies provides an estimate of the maximal imputation performance for the out-panel samples (Fig. 8). Even at 1x depth, PPVs reached 0.86 to 0.87 for deletions with 84 to 85% recall (Fig. 8a, b). At 4x depth, the highest PPV of 0.92 was recovered for deletions, with 93% recall using non-logical filtering (Fig. 8a, b). Imputed duplications for the out-panel samples reached a much lower PPV than deletions (Fig. 8a), but were consistent with the PPV values of duplications in in-panel samples (Fig. 4).

**Figure 8.**
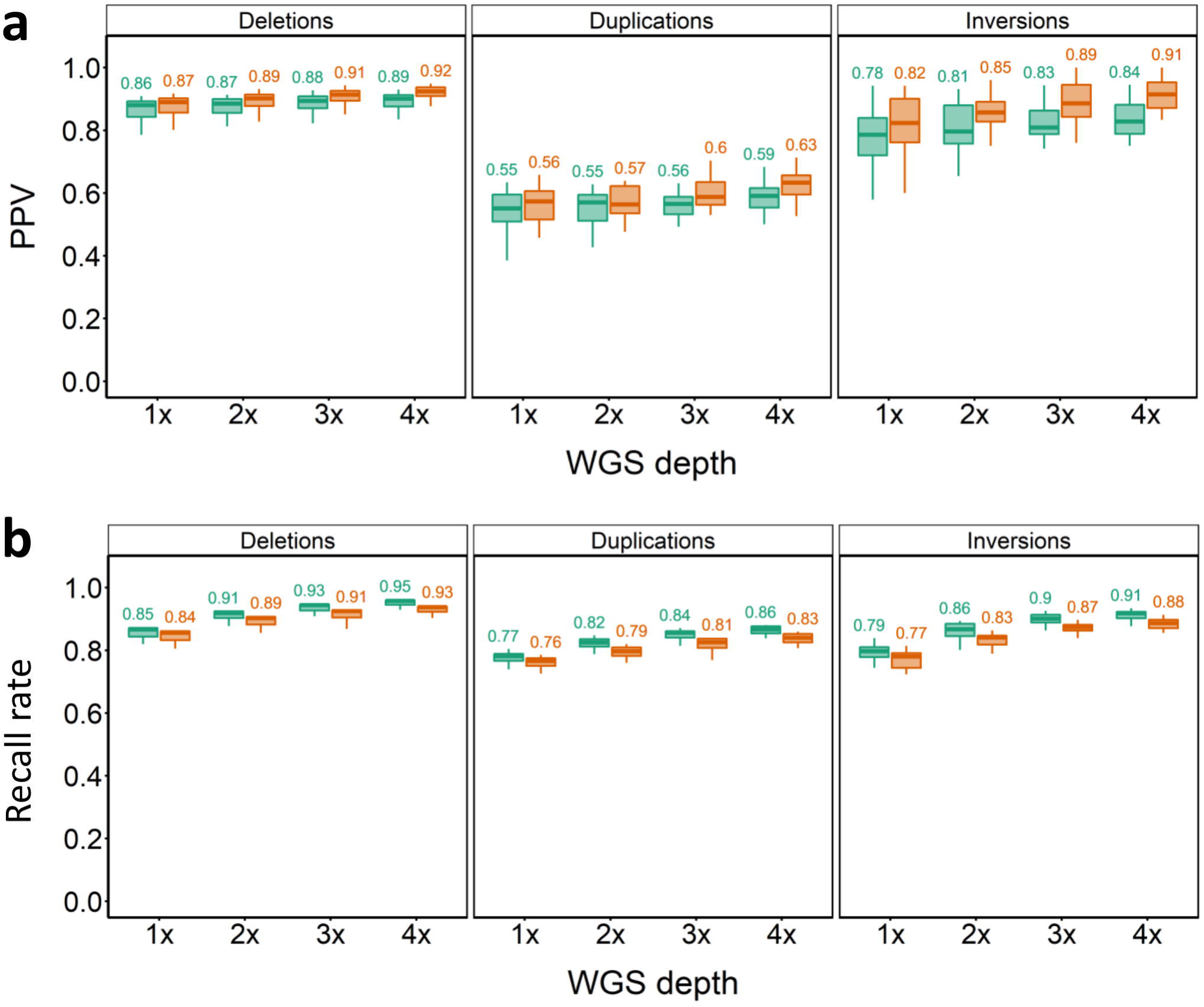
Merged SV imputation results for out-panel samples. PPV (**a**) and recall rate (**b**) for SV imputation across different SV classes, WGS depths and strategies at a PP cut-off of 0.9. Green plots indicate PPVs for the ‘logical’ and brown for ‘non-logical’ SV filtering approaches (see Methods).

## Discussion

It has been argued that genotype imputation using low-coverage WGS against a reference panel is poised to bring about a paradigm shift in genetics by enabling genome-wide variant analyses in a highly cost-effective manner [12]. Beyond human genetics, the farmed animal community has begun to uptake this strategy, for example a recent study of cattle that benchmarked different imputation methods [47]. Our study demonstrates the successful application of GLIMPSE for genome-wide variant imputation using low coverage WGS in Atlantic salmon. We advance on previous work by capturing a global set of gold-standard SVs [29] in the reference panel. While imputation performance was lower for SVs than for SNVs, this must be framed against the challenge of accurately calling SVs directly using WGS data, which suffers from extremely high false positive rates, approaching 90% in Atlantic salmon, demanding extensive curation and filtering [29]. In this context, the accuracy and recall of SV calls achieved using low coverage WGS was impressive, for instance reaching 92% accuracy and 94% recall for deletions with 4x data. Moreover, the pipeline developed in this study provides a broadly applicable framework for the simultaneous imputation of SVs and SNVs across other livestock species, provided high-quality reference panels are available. A caveat of our method is that only known high-confidence SVs defined within the reference panel are imputed. This is well-illustrated by the out-panel samples tested, for which approximately half the high-confidence SVs detected *de novo* were not present in the reference panel. This limitation requires updating the SV reference panel to ensure all variants of interest are captured in a target population (discussed further below). Interestingly, the equivalent imputation performance of the Hendrix-Genetics Landcatch strain (i.e. out-panel) samples to the in-panel samples indicates that these farmed fish share extensive haplotypes with the wild reference panel, with the caveat that the sample size was limited to n=20. This highlights the applicability of combined genome-wide SV and SNV imputation with low-coverage WGS in commercial Atlantic salmon breeding programmes.

An interesting observation when performing SV imputation with GLIMPSE was that the supplementation of SV GLs (i.e. GLIMPSE+SVtyper) increased imputation accuracy compared to imputation based solely on LD with SNVs in the same haplotypes. The impact of including SV GLs on SV imputation accuracy was most pronounced for longer SVs of all classes, even when using 1x depth. By combining the unified SV calls from both strategies, we achieved the best combination of accuracy and recall, at the cost of a more complex computational workflow. However, notable gains in performance using GLIMPSE+SVtyper required 3x to 4x depths, which adds further sequencing costs compared to relying purely on LD with SNVs. We also observed differences in imputation of different SV classes, with performance for deletions exceeding that for duplications and inversions. Duplications were the worst performing SV class, with mean PPV never exceeding ∼60%, although markedly higher PPVs were achieved for shorter duplications. Differences in imputation performance across different type of SVs can arise for many reasons, including different allele frequency distributions and perhaps most importantly, the comparative ability to call these variants accurately from short-read sequencing data for both the reference panels and individual low coverage sequencing data. In this respect, the relative performance of imputation across different SV classes closely matched differences in the accuracy of the original reference panel used for the same classes of SV [29]. Another notable observation was similar imputation performance for individuals sampled from North and South Norwegian rivers. Although we anticipated differences in performance between these two sample sets, the nearly identical results may be attributed to the inclusion of individuals from various rivers in both the North and South Norwegian test datasets.

Our study has several obvious limitations and the method we report can be improved in the future. Firstly, our reference panel was based on mapping WGS reads to the previous Atlantic salmon reference genome ICSAG_v2 [23], which was recently superseded by higher quality assemblies for multiple phylogeographic lineages [24,25]. A genome for a Norwegian fish represents the current Atlantic salmon reference (Ssal_v3.1) and has been annotated by NCBI (accession: GCF_905237065) and Ensembl (https://www.ensembl.org/Salmo_salar/). In the future, a new reference panel can be built taking advantage of these assemblies, which have much greater contiguity and completeness and fewer errors than ICSAG_v2, owing to their construction with long-read technologies (e.g. [24]). The Ssal_v3.1 genome has been annotated using extensive functional genomic data (RNA-Seq, ChIP-Seq, ATAC-Seq) generated across development and tissue types by the European AQUA-FAANG consortium (https://www.aqua-faang.eu/) [48]. A future reference panel updated to Ssal_v3.1 could cross-reference imputed variants with this rich functional annotation, including regulatory elements in non-coding regions that control gene expression, enhancing scope to identify candidate causal variants in QTL regions.

Another way to improve our reference panel would be to incorporate a broader definition of genetic diversity for Atlantic salmon. As our reference panel is comprised solely of European (mainly Norwegian) Atlantic salmon populations, it should be updated in the future to capture other lineages. Atlantic salmon is split into two major phylogeographic lineages that separated more than 1 million years ago, with limited subsequent gene flow [49], leading to major genomic differences and an argument for two ‘sub-species’ [25,50]. Wild North American salmon have been used as separate founders for aquaculture strains [28] and are of substantial interest in terms of molecular ecology and applied conservation genetics. Due to the significant evolutionary distance involved, the performance of our reference panel with low-coverage WGS data from North American salmon will likely be substantially reduced. As WGS data is available for North American salmon [28], future updates to the reference panel should capture this, as well as other available genetic variation representing the major Atlantic salmon lineages. An improved reference panel that includes more individuals sequenced at higher depths should help capture additional genomic variants that could be missing or underrepresented in the current reference panel.

Another limitation of our reference panel is that SVs were defined solely by short-read sequencing [29]. The current state of the art for SV discovery is long-read sequencing, which enables the discovery of more types of SVs compared to short-read WGS and with reduced false positive calls [4]. Moreover, the method used previously by Bertolotti et al. [29] for SV discovery excluded insertion variants. Consequently, our current reference panel is likely far from comprehensive of the true catalogue of SVs in Atlantic salmon genomes. There are several ways to enhance representation of SVs in a future Atlantic salmon reference panel. The lack of insertions could be addressed using an SV calling algorithm that is specifically designed to capture this class of variant [51]. More importantly, long-read sequencing should be leveraged to characterise SVs more comprehensively, although this currently remains cost prohibitive for large-scale population genetics. Given the availability of multiple high-quality assemblies for different Atlantic salmon lineages built with long-read data [24], there is an exciting opportunity to produce a graph-genome (alternatively called pan-genome) representation of genetic variation in the Atlantic salmon genome, which could be used for subsequent genotyping with short-read WGS [52]. This strategy was performed recently in a related salmonid species (whitefish; *Coregonus* spp.) to define a much larger set of SVs than what was captured with short-read data [53]. Graph/pan genome representations also allow variants to be captured within regions of the genome that are absent in a single reference genome [54]. While graph genomes have not yet been used alongside imputation with low-coverage WGS in the published literature, this will be a valuable strategy in the future to capture a comprehensive spectrum of SVs and explore their impact on phenotypic variation.

## Conclusions

We demonstrate the promise of leveraging a reference panel to call SNVs and SVs with low-coverage WGS. The strategy we developed is generic and can be readily transferred to other species with a high-quality set of SNVs and SVs defined in a representative panel of individuals, providing opportunities to enhance the resolution of GWAS in a cost-effective manner by capturing a wider spectrum of potential causal variants.

## Declarations

**Ethics approval and consent to participate:** Not applicable

**Consent for publication:** Not applicable

**Availability of data and materials**: The twenty WGS samples generated for the out-panel samples are available through NCBI (BioProject accession - PRJNA917857). All WGS and genetic variant data associated with the reference panel, including the in-panel WGS data, is available in public archives, as described by Bertolotti et al. (2020) [29] and Sinclair-Waters et al. (2022) [30].

## Authors’ contributions

Designed study: MKG, JGDP, and DJM. Secured funding: RDH and DJM. Provided out-panel samples used for WGS: AH. Performed imputation analyses: MKG and JGDP. Performed detection of SVs and SNVs in out-panel samples: MKG. Supported data analysis: DR and JGDP. Prepared figures: MKG. Drafted manuscript: MKG and DJM. Provided intellectual inputs, and contributed to data interpretation, writing and finalization of manuscript: All authors. All authors read and approved the final manuscript

## Acknowledgements

Prof. Sigbjørn Lien and Dr Torfinn Nome (Norwegian University of Life Sciences) shared the SNV data from Sinclair-Waters et al. 2022.

## Additional files

### Additional file 1 Table S1

File format: XLSX

Title: Details of the samples in the reference panel

### Additional file 1 Table S2

File format: XLSX

Title: Details of the In-panel samples used in the study.

### Additional file 1 Table S3

File format: XLSX

Title: Details of the Out-panel samples used in the study.

### Additional file 1 Table S4

File format: XLSX

Title: Number of SNVs imputed at various PPs and sequencing depths.

### Additional file 2 Figure S1

File format: DOC

Title: SNV imputation results for Southern Norway samples (n=49).

PPV (a) and recall rate (b) for SNVs across tested WGS depths and PP cut-offs.

### Additional file 2 Figure S2

File format: DOC

Title: SNV imputation results for Northern Norway samples (n=49).

PPV (a) and recall rate (b) for SNVs across tested WGS depths and PP cut-offs.

### Additional file 2 Figure S3

File format: DOC

Title: Performance of Southern Norway sample (n=49) deletion SV imputation.

Description: PPV (a) and recall rate (b) for across different PP cut-offs, sequencing depths and strategies used. Red and blue plots indicate PPVs when the pipeline was supplied with SNP GLs only, or both SNP and SV GLs, respectively.

### Additional file 2 Figure S4

File format: DOC

Title: Performance of Northern Norway sample (n=49) deletion SV imputation.

Description: PPV (a) and recall rate (b) for across different PP cut-offs, sequencing depths and strategies used. Red and blue plots indicate PPVs when the pipeline was supplied with SNP GLs only, or both SNP and SV GLs, respectively.

### Additional file 2 Figure S5

File format: DOC

Title: Performance of in-panel sample (n=98) duplication SV imputation.

Description: PPV (a) and recall rate (b) for across different PP cut-offs, sequencing depths and strategies used. Red and blue plots indicate PPVs when the pipeline was supplied with SNP GLs only, or both SNP and SV GLs, respectively.

### Additional file 2 Figure S6

File format: DOC

Title: Performance of in-panel sample (n=98) inversion SV imputation.

Description: PPV (a) and recall rate (b) for across different PP cut-offs, sequencing depths and strategies used. Red and blue plots indicate PPVs when the pipeline was supplied with SNP GLs only, or both SNP and SV GLs, respectively.

### Additional file 2 Figure S7

File format: DOC

Title: Performance of Southern Norway sample (n=49) duplication SV imputation.

Description: PPV (a) and recall rate (b) for across different PP cut-offs, sequencing depths and strategies used. Red and blue plots indicate PPVs when the pipeline was supplied with SNP GLs only, or both SNP and SV GLs, respectively.

### Additional file 2 Figure S8

File format: DOC

Title: Performance of Northern Norway sample (n=49) duplication SV imputation.

Description: PPV (a) and recall rate (b) for across different PP cut-offs, sequencing depths and strategies used. Red and blue plots indicate PPVs when the pipeline was supplied with SNP GLs only, or both SNP and SV GLs, respectively.

### Additional file 2 Figure S9

File format: DOC

Title: Performance of Southern Norway sample (n=49) inversion SV imputation.

Description: PPV (a) and recall rate (b) for across different PP cut-offs, sequencing depths and strategies used. Red and blue plots indicate PPVs when the pipeline was supplied with SNP GLs only, or both SNP and SV GLs, respectively.

### Additional file 2 Figure S10

File format: DOC

Title: Performance of Northern Norway sample (n=49) inversion SV imputation.

Description: PPV (a) and recall rate (b) for across different PP cut-offs, sequencing depths and strategies used. Red and blue plots indicate PPVs when the pipeline was supplied with SNP GLs only, or both SNP and SV GLs, respectively.

### Additional file 2 Figure S11

File format: DOC

Title: Impact of SV length on imputation accuracy of in-panel samples (N=98).

PPVs for SV imputation at different SV length cut-offs are shown. The x-axis of each plot represents the length bins into which SVs were classified. Blue plots represent WGS depth of 1x, and green plots represent WGS depth of 4x. DEL-PPV, DUP-PPV, and INV-PPV correspond to the PPVs for deletions, duplications, and inversions, respectively. The numbers above each box plot indicate the mean PPV, while the numbers in red below the boxplots represent the total number of SVs in each length category.

### Additional file 2 Figure S12

File format: DOC

Title: Merged SV imputation results for Southern Norway samples (n=49).

PPV (a) and recall rate (b) for SV imputation across different SV classes, WGS depths and strategies at a PP cut-off of 0.9. Green plots indicate PPVs for the ‘logical’ and brown for ‘non-logical’ SV filtering approaches (see Methods).

### Additional file 2 Figure S13

File format: DOC

Title: Merged SV imputation results for Northern Norway samples (n=49).

PPV (a) and recall rate (b) for SV imputation across different SV classes, WGS depths and strategies at a PP cut-off of 0.9. Green plots indicate PPVs for the ‘logical’ and brown for ‘non-logical’ SV filtering approaches (see Methods).

### Additional file 2 Figure S14

File format: DOC

Title: Allele frequency distribution of overlapping and non-overlapping SVs.

Description: Allele frequency distributions are shown for overlapping (g) and non-overlapping (h) homozygous (i.e. 1/1) SVs, as well as for overlapping (i) and non-overlapping (j) heterozygous (0/1) SVs, in the out-panel samples.

### Additional file 2 Figure S15

File format: DOC

Title: Performance of out-panel sample (n=20) duplication SV imputation.

Description: PPV (a) and recall rate (b) for across different PP cut-offs, sequencing depths and strategies used. Red and blue plots indicate PPVs when the pipeline was supplied with SNP GLs only, or both SNP and SV GLs, respectively.

### Additional file 2 Figure S16

File format: DOC

Title: Performance of out-panel sample (n=20) inversion SV imputation.

Description: PPV (a) and recall rate (b) for across different PP cut-offs, sequencing depths and strategies used. Red and blue plots indicate PPVs when the pipeline was supplied with SNP GLs only, or both SNP and SV GLs, respectively.

