## supplementary fig s1 to s16 for "High performance imputation of structural and single nucleotide variants using low-coverage whole genome sequencing"

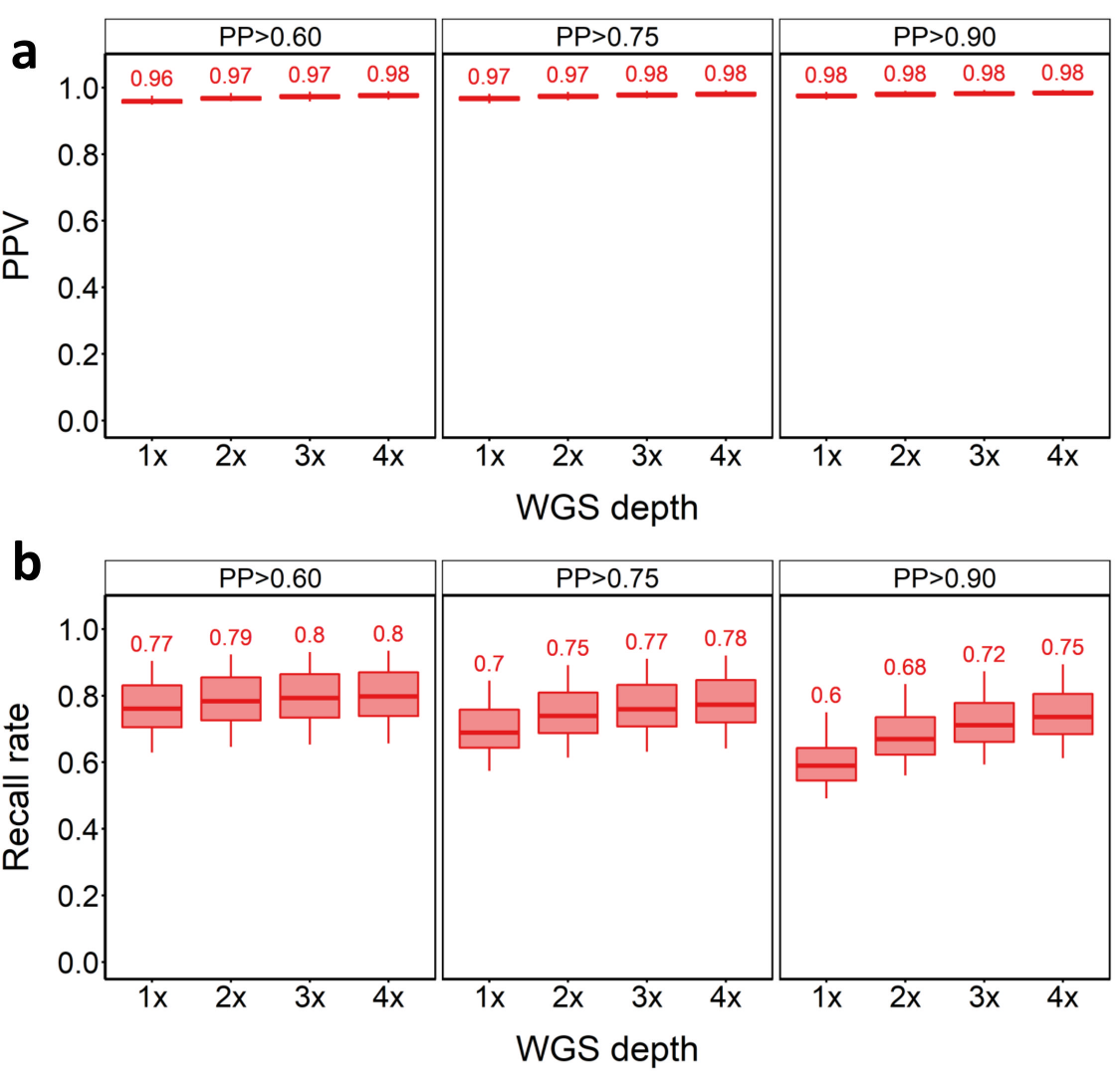
**Supplementary Figures**

**Supplementary Figure 1. SNV imputation results for Southern Norway samples (n=49).**

PPV (a) and recall rate (b) for SNVs across tested WGS depths and PP cut-offs.

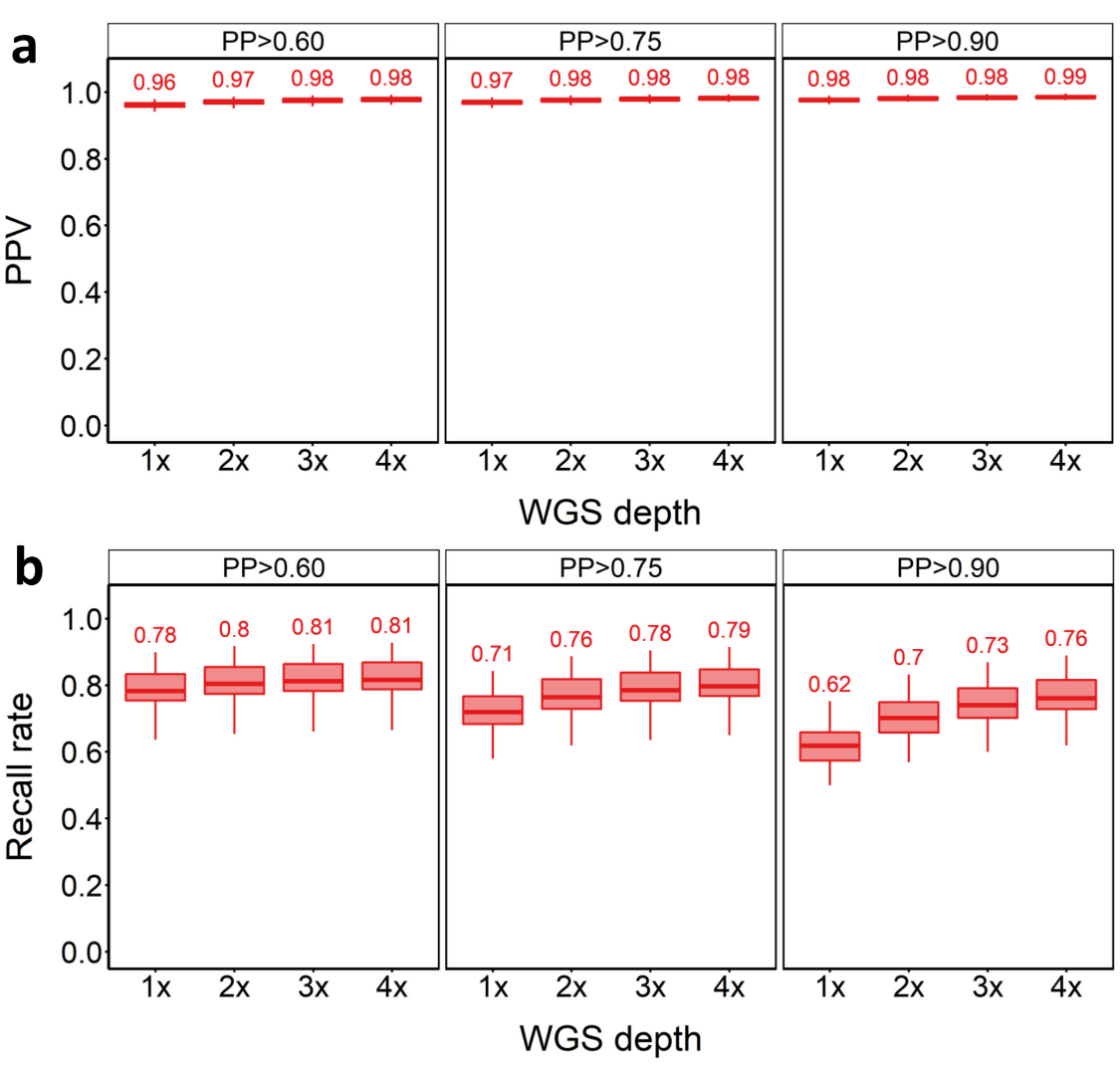

**Supplementary Figure 2. SNV imputation results for Northern Norway samples (n=49).**

PPV (a) and recall rate (b) for SNVs across tested WGS depths and PP cut-offs.

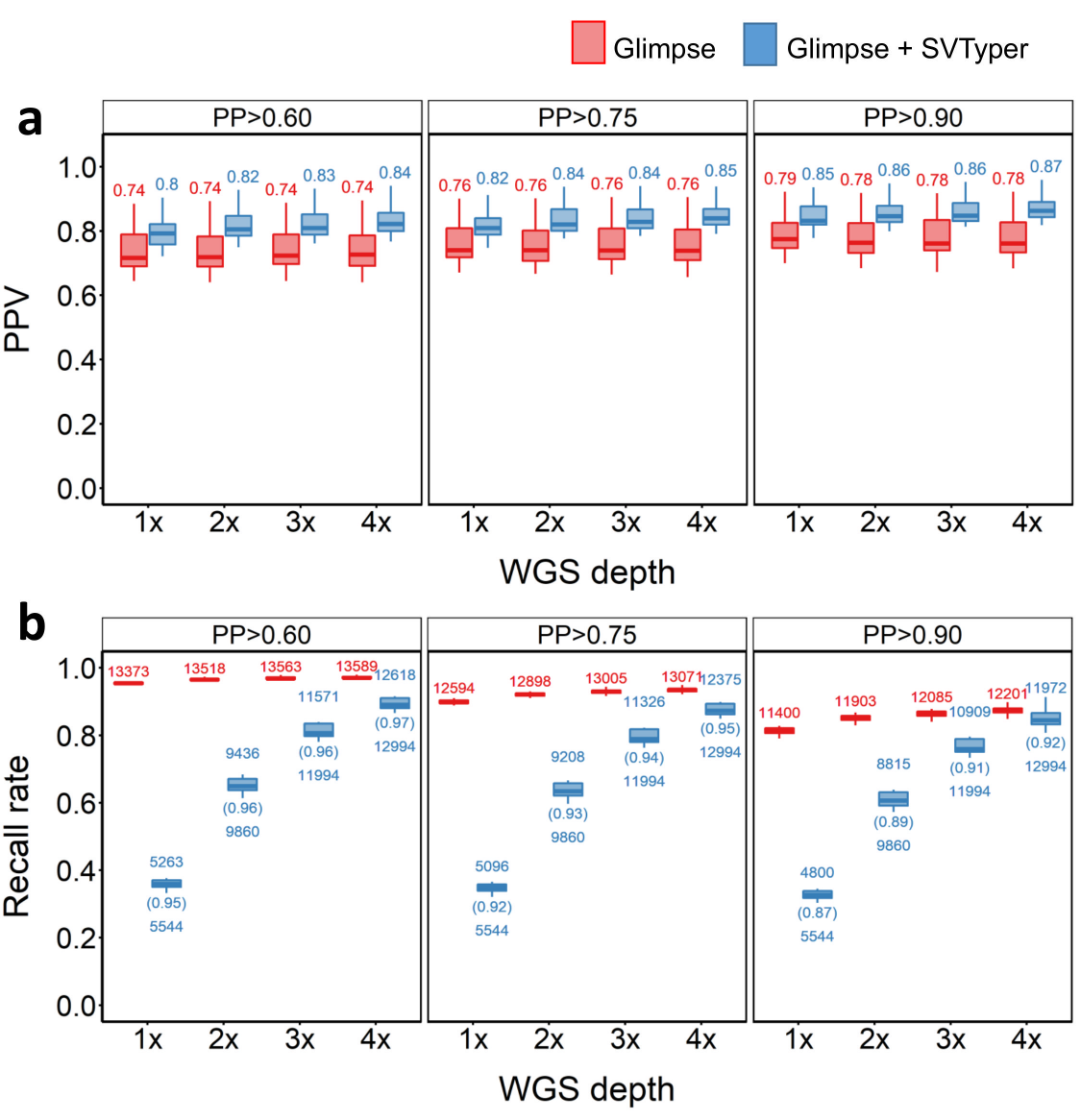

**Supplementary Figure 3.**  **Performance of Southern Norway sample (n=49) deletion SV imputation.**

PPV (**a**) and recall rate (**b**) for across different PP cut-offs, sequencing depths and strategies used. Red and blue plots indicate PPVs when the pipeline was supplied with SNP GLs only, or both SNP and SV GLs, respectively.

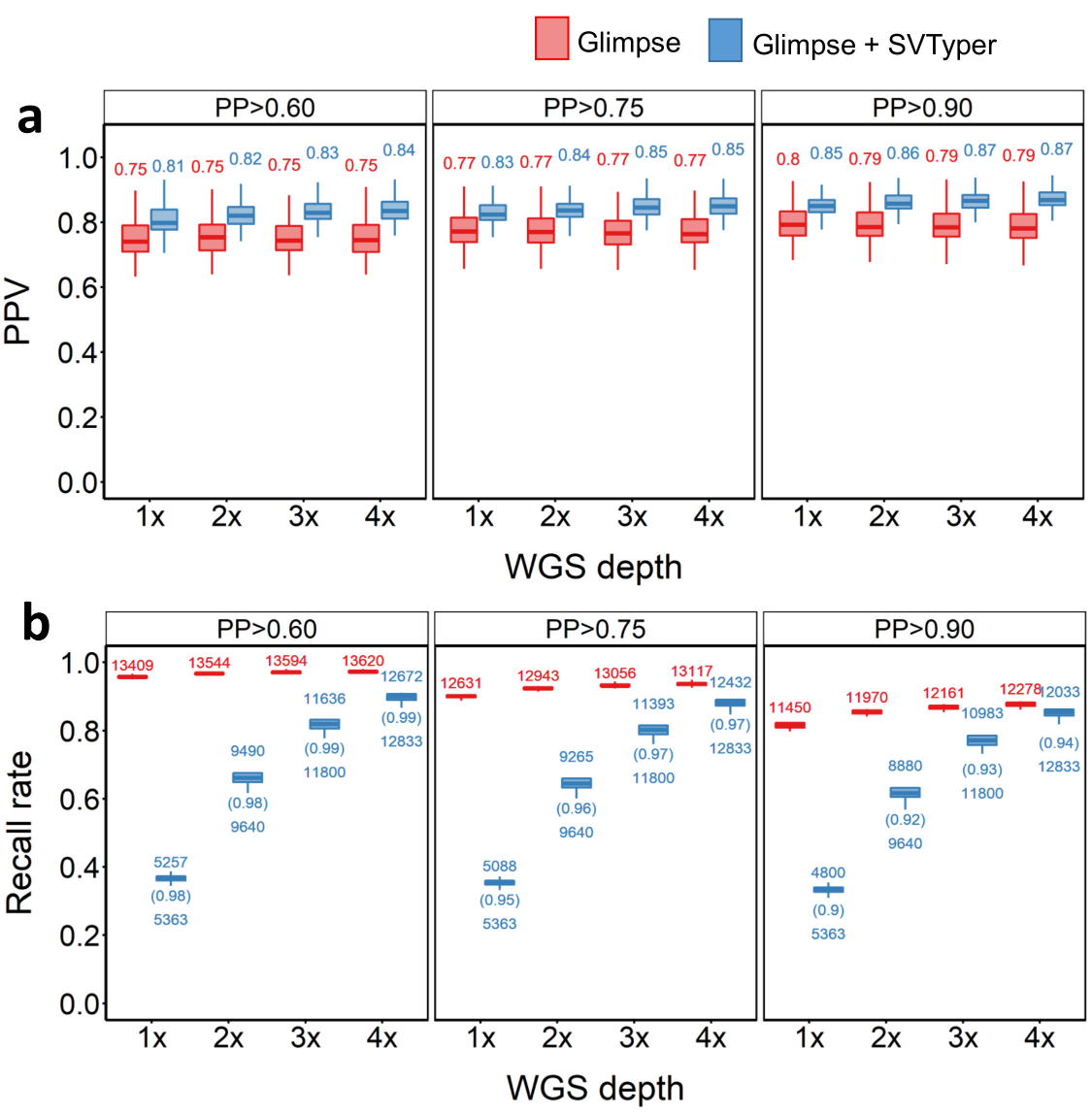

**Supplementary Figure 4.**  **Performance of Northern Norway sample (n=49) deletion SV imputation.**

PPV (**a**) and recall rate (**b**) for across different PP cut-offs, sequencing depths and strategies used. Red and blue plots indicate PPVs when the pipeline was supplied with SNP GLs only, or both SNP and SV GLs, respectively.

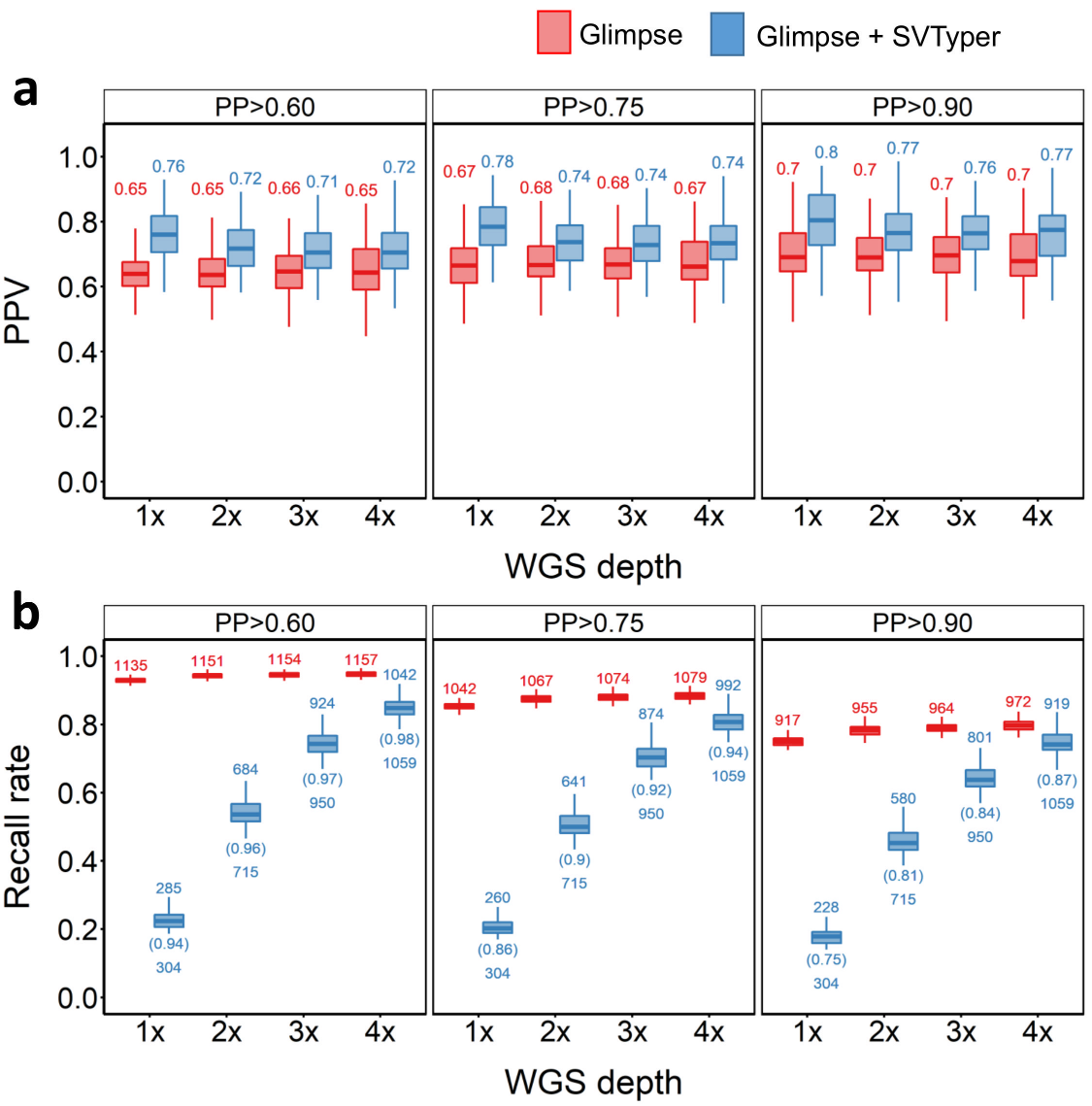

**Supplementary Figure 5. Performance of in-panel sample (n=98) duplication SV imputation.**

Description: PPV (**a**) and recall rate (**b**) for across different PP cut-offs, sequencing depths and strategies used. Red and blue plots indicate PPVs when the pipeline was supplied with SNP GLs only, or both SNP and SV GLs, respectively.

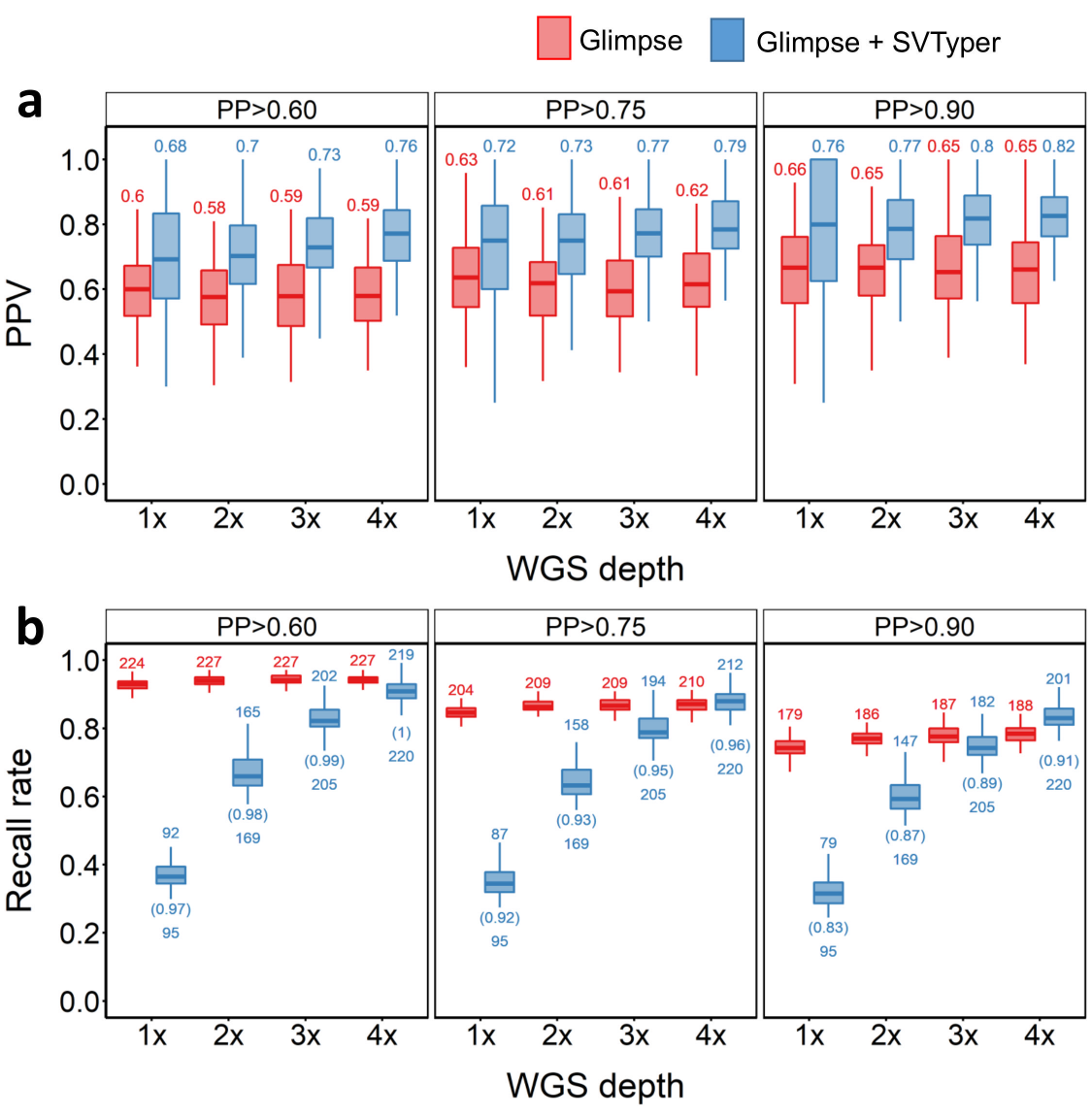

**Supplementary Figure 6. Performance of in-panel sample (n=98) inversion SV imputation.**

Description: PPV (**a**) and recall rate (**b**) for across different PP cut-offs, sequencing depths and strategies used. Red and blue plots indicate PPVs when the pipeline was supplied with SNP GLs only, or both SNP and SV GLs, respectively.

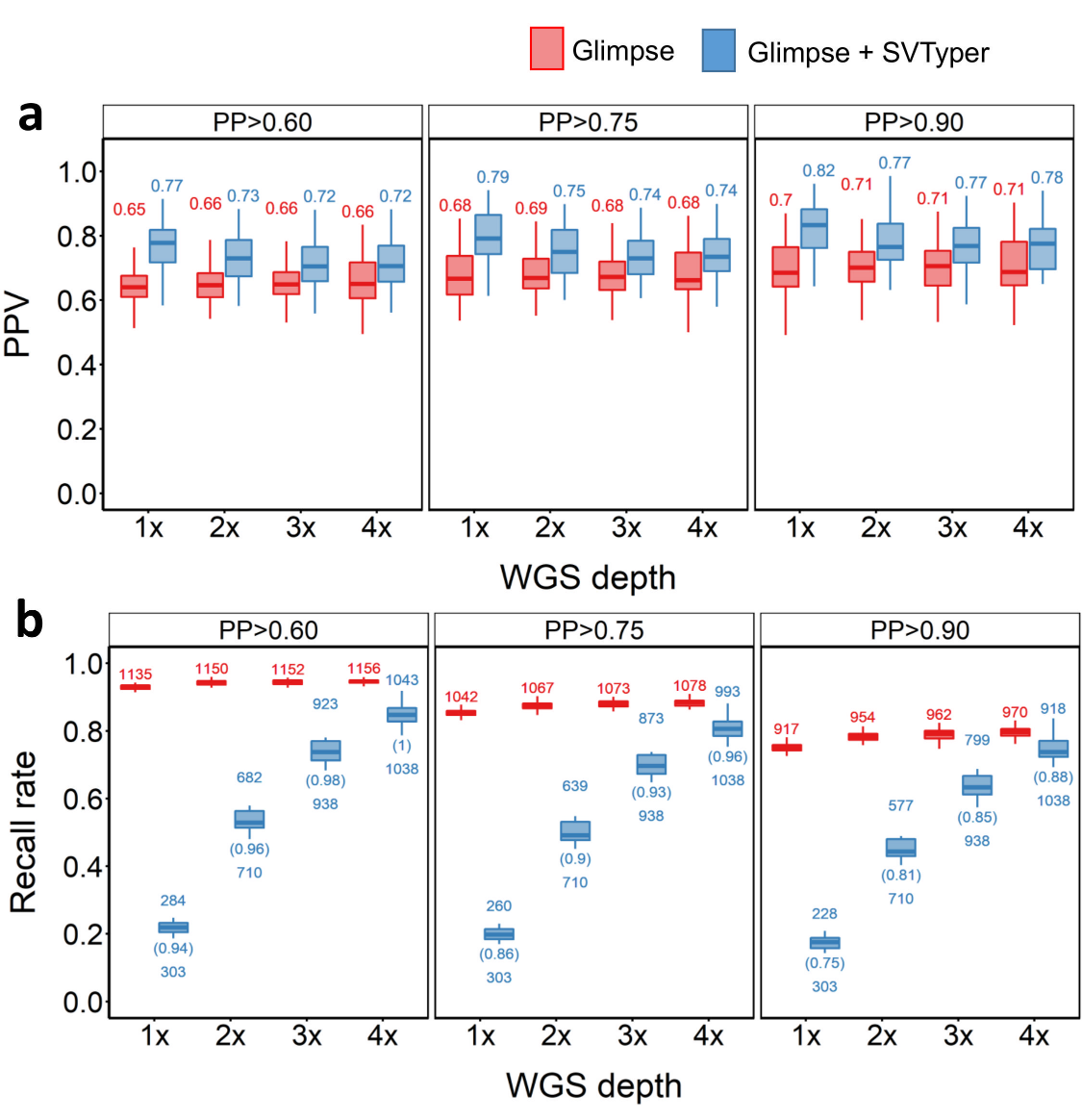

**Supplementary Figure 7. Performance of Southern Norway sample (n=49) duplication SV imputation.**

PPV (a) and recall rate (b) for across different PP cut-offs, sequencing depths and strategies used. Red and blue plots indicate PPVs when the pipeline was supplied with SNP GLs only, or both SNP and SV GLs, respectively.

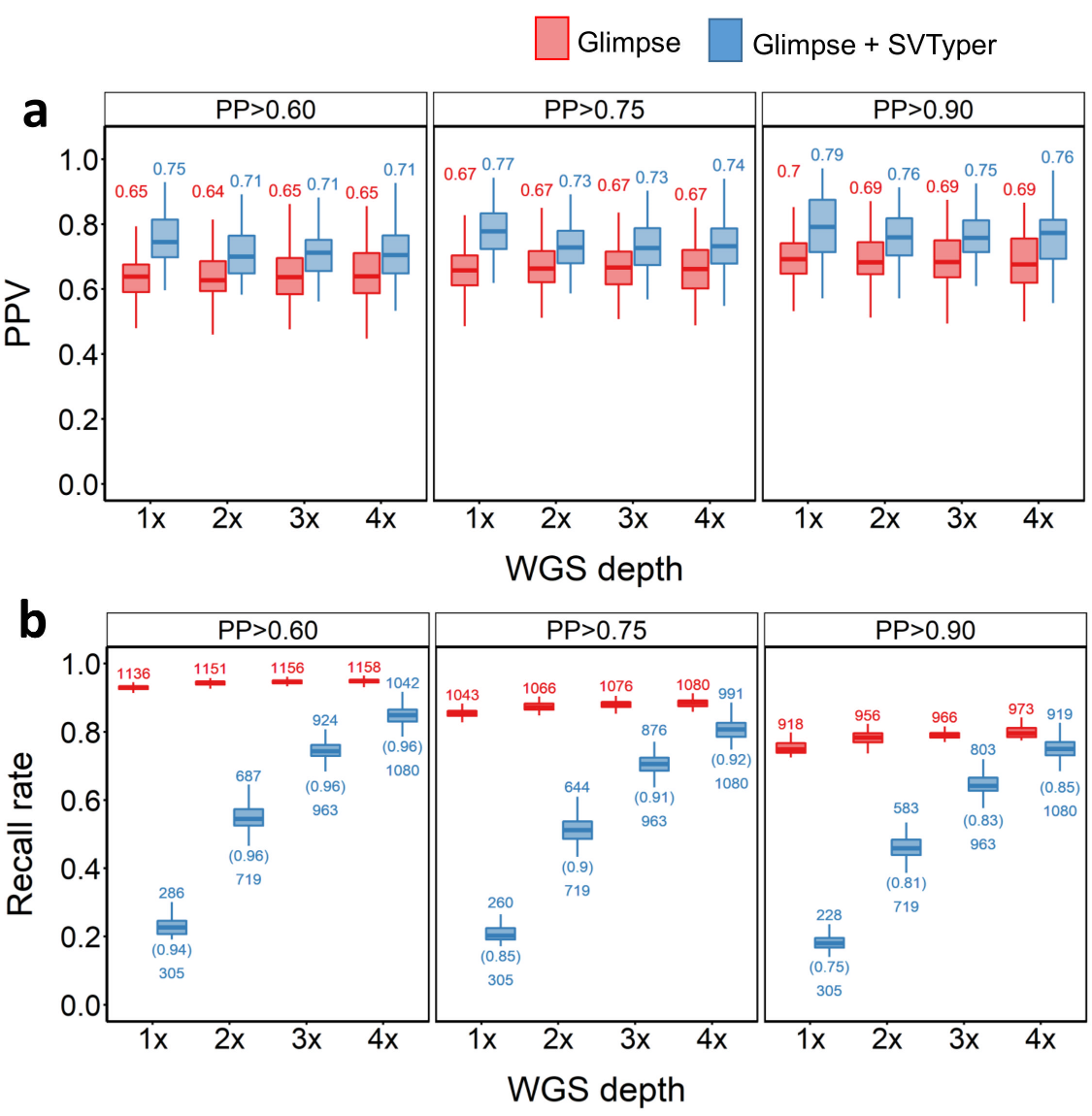

**Supplementary Figure 8.**  **Performance of Northern Norway sample (n=49) duplication SV imputation.**

PPV (**a**) and recall rate (**b**) for across different PP cut-offs, sequencing depths and strategies used. Red and blue plots indicate PPVs when the pipeline was supplied with SNP GLs only, or both SNP and SV GLs, respectively.

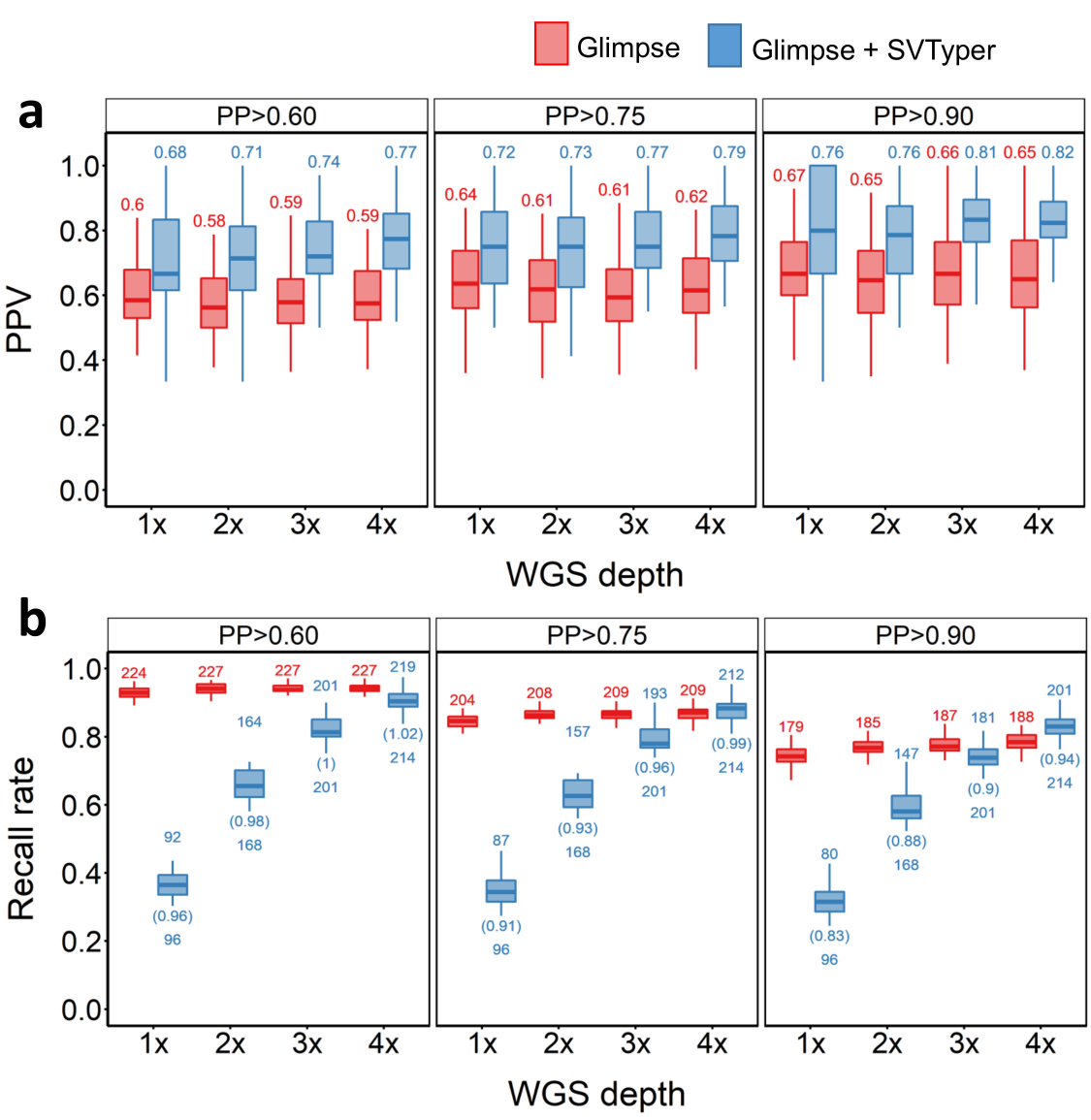

**Supplementary Figure 9. Performance of Southern Norway sample (n=49) inversion SV imputation.**

PPV (a) and recall rate (b) for across different PP cut-offs, sequencing depths and strategies used. Red and blue plots indicate PPVs when the pipeline was supplied with SNP GLs only, or both SNP and SV GLs, respectively.

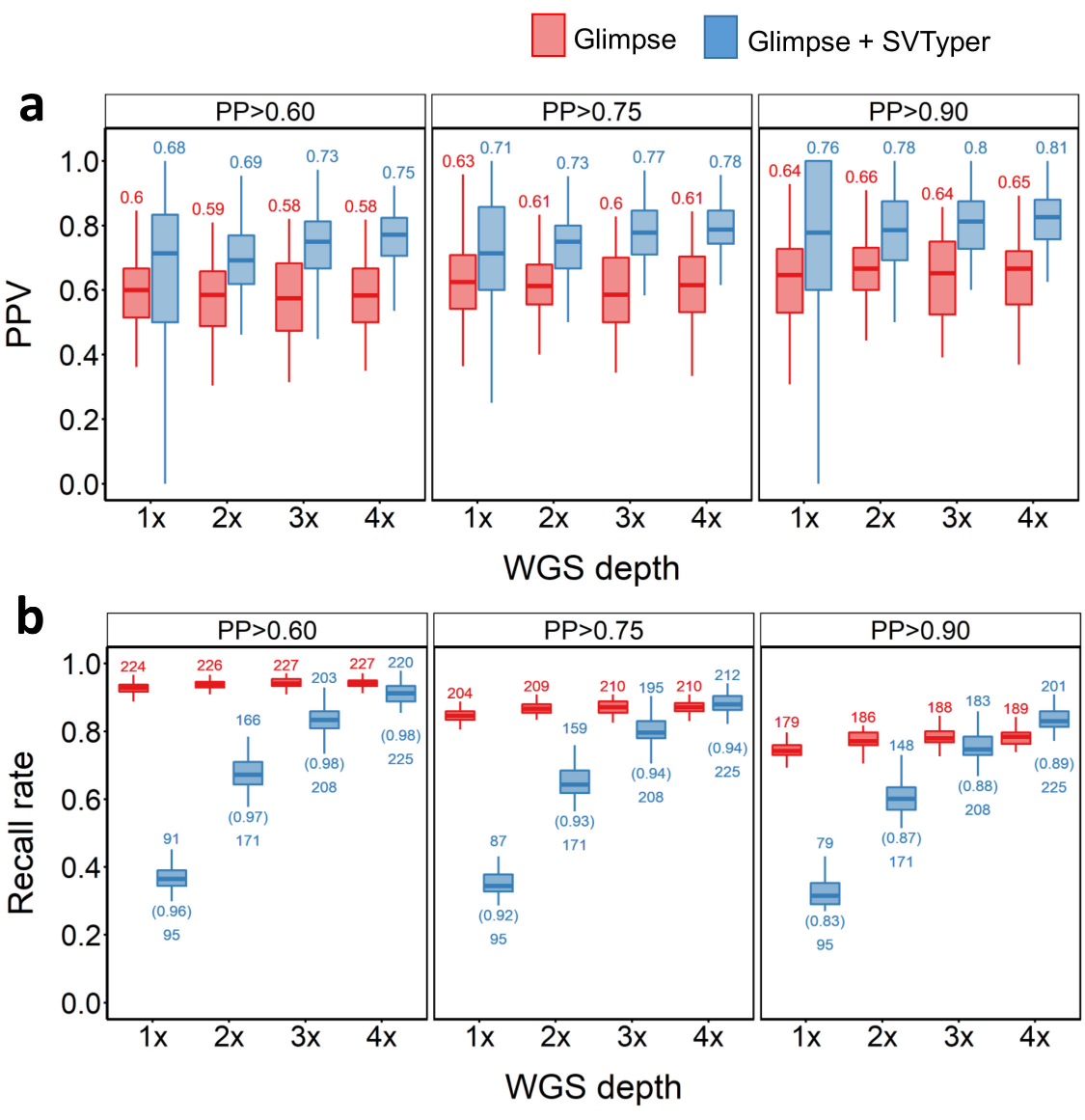

**Supplementary Figure 10.**  **Performance of Northern Norway sample (n=49) inversion SV imputation.**

PPV (**a**) and recall rate (**b**) for across different PP cut-offs, sequencing depths and strategies used. Red and blue plots indicate PPVs when the pipeline was supplied with SNP GLs only, or both SNP and SV GLs, respectively.

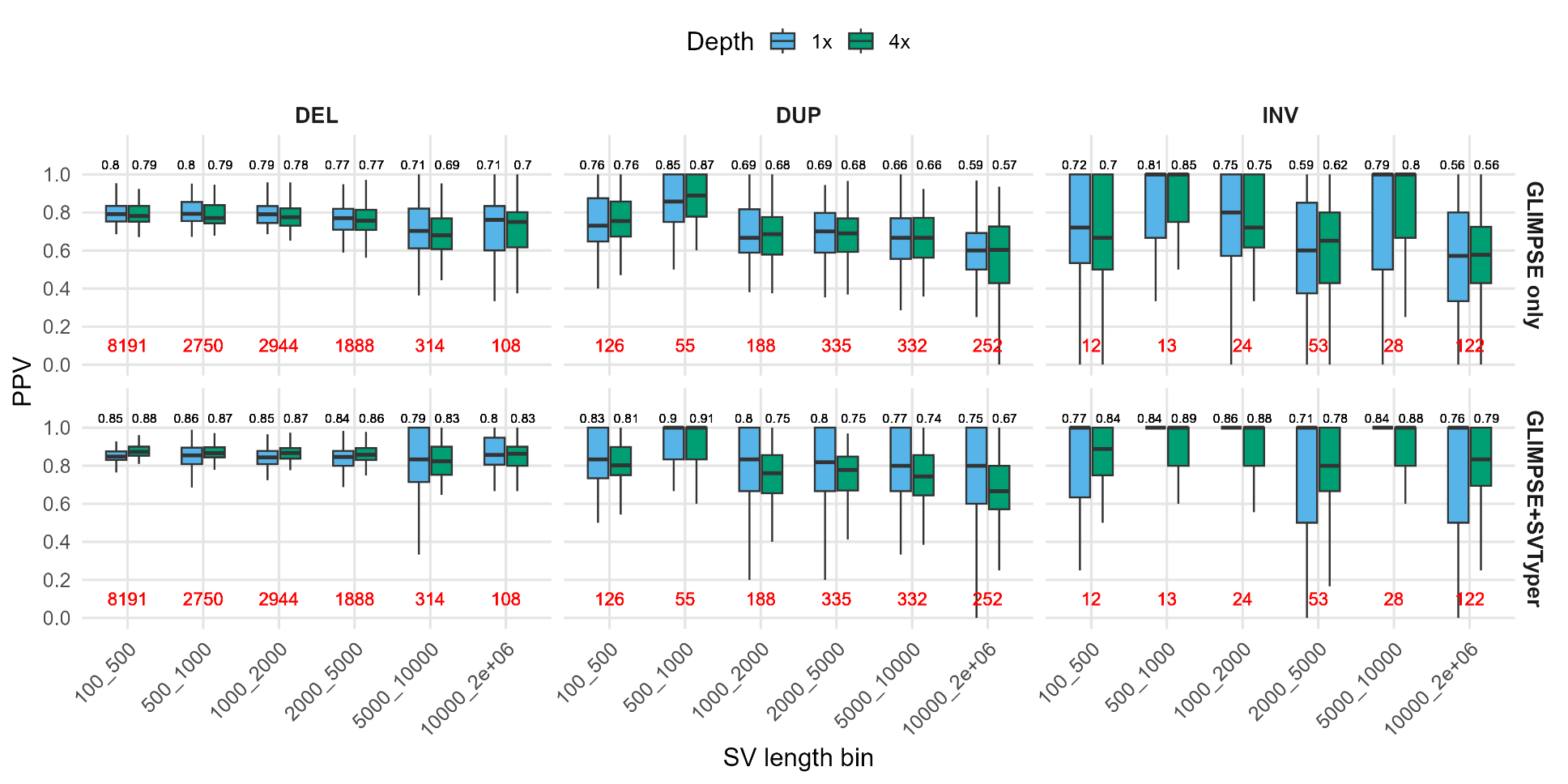
**Supplementary Figure 11. Impact of SV length on SV imputation accuracy of in-panel samples (n=98)**

PPVs for different SV length bins are shown at 1x and 4x depths. DEL, DUP, and INV are deletions, duplications, and inversions, respectively. Numbers above boxplots indicate mean PPV, while red numbers below boxplots describe the number of SVs per length bin.

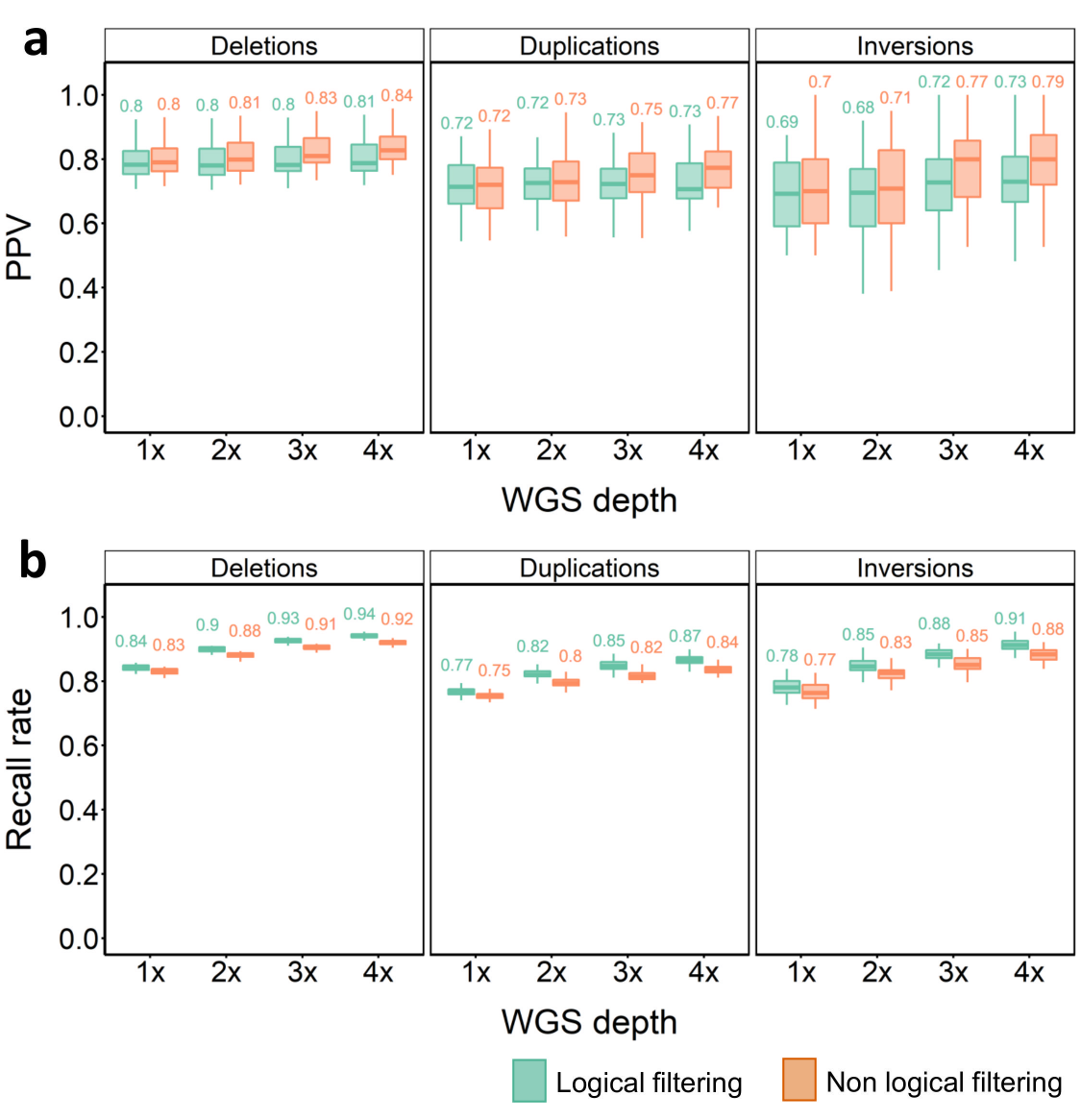

**Supplementary Figure 12. Merged SV imputation results for Southern Norway samples.**

PPV (**a**) and recall rate (**b**) for SV imputation across different SV classes, WGS depths and strategies at a PP cut-off of 0.9. Green plots indicate PPVs for the ‘logical’ and brown for ‘non-logical’ SV filtering approaches (see Methods).

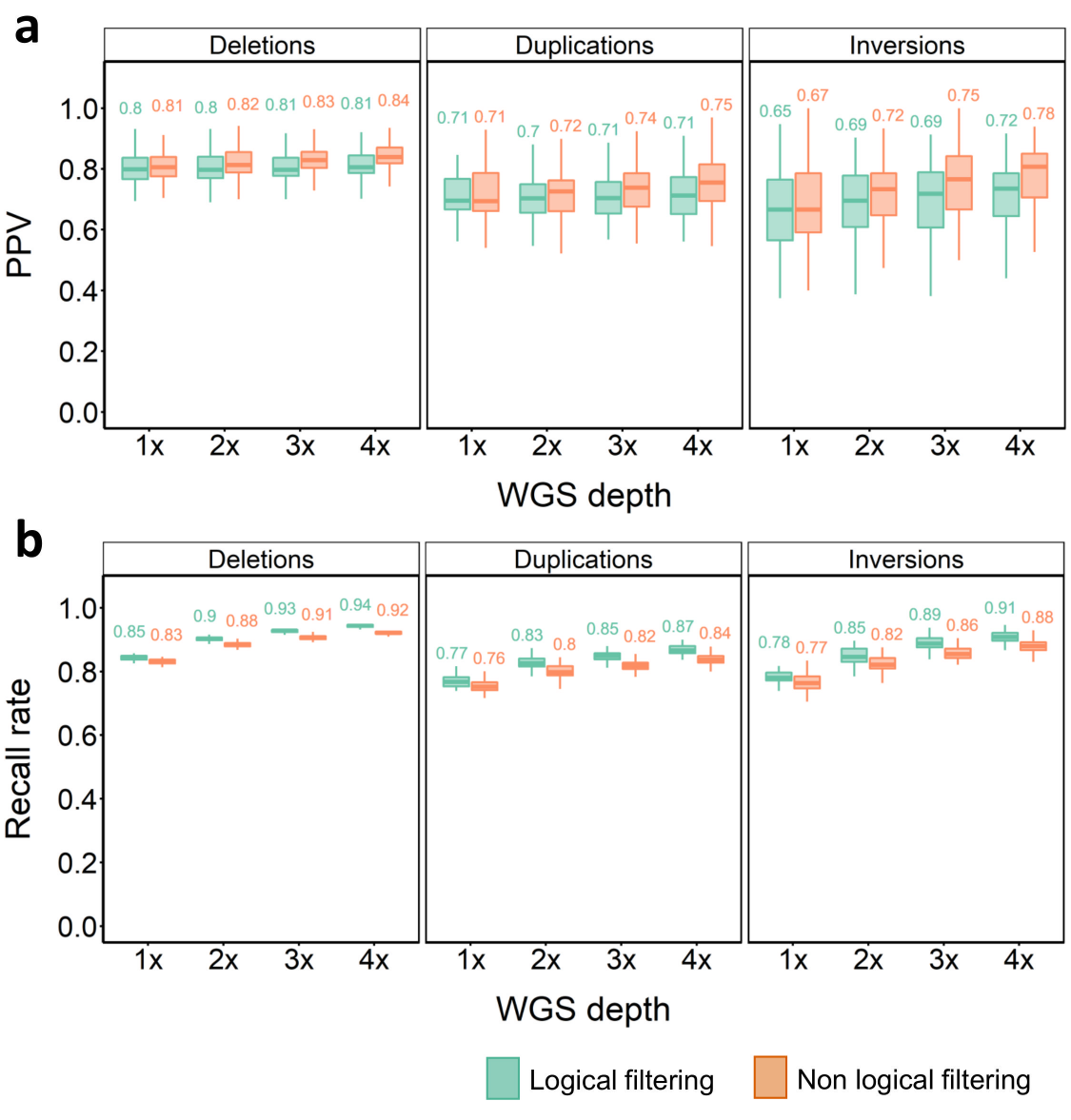

**Supplementary Figure 13. Merged SV imputation results for Northern Norway samples.**

PPV (**a**) and recall rate (**b**) for SV imputation across different SV classes, WGS depths and strategies at a PP cut-off of 0.9. Green plots indicate PPVs for the ‘logical’ and brown for ‘non-logical’ SV filtering approaches (see Methods).

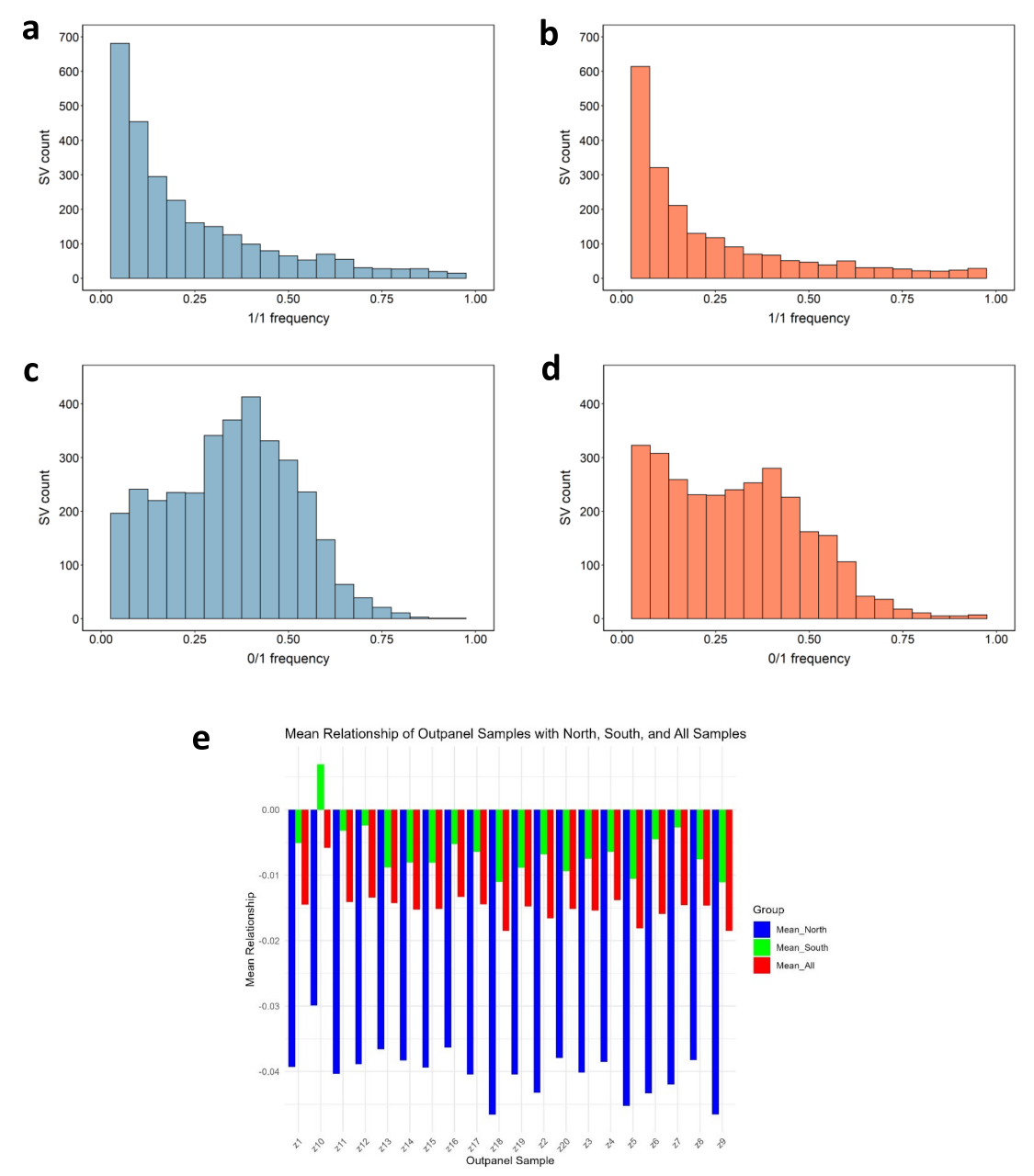

**Supplementary Figure 14. Allele frequency distribution of overlapping and non-overlapping SVs.**

Allele frequency distributions are shown for overlapping (**a**) and non-overlapping (**b**) homozygous (i.e. 1/1) SVs, as well as for overlapping (**c**) and non-overlapping (**d**) heterozygous (0/1) SVs, in the out-panel samples.

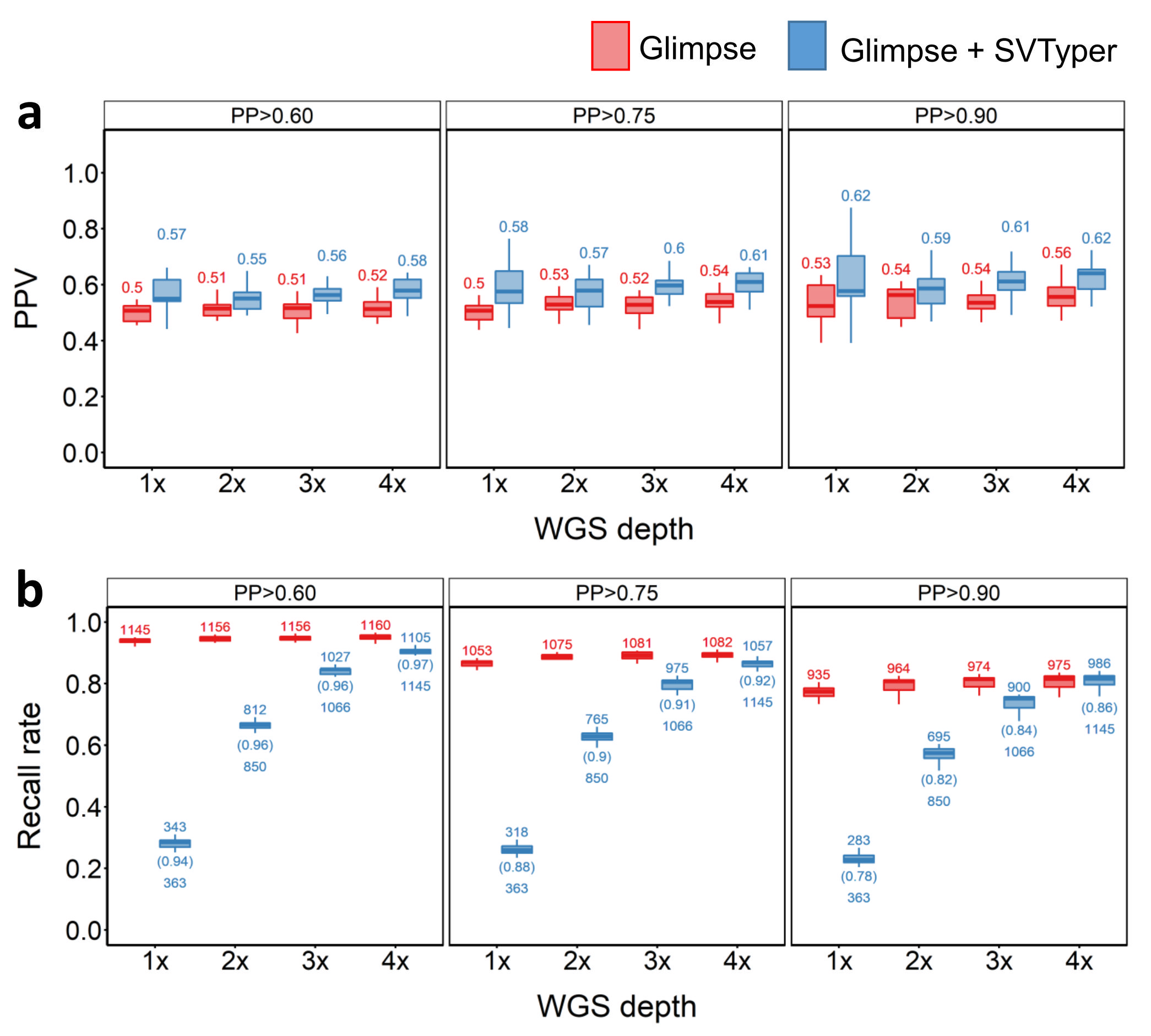

**Supplementary Figure 15. Performance of out-panel sample (n=20) duplication SV imputation.** PPV (**a**) and recall rate (**b**) for across different PP cut-offs, sequencing depths and strategies used. Red and blue plots indicate PPVs when the pipeline was supplied with SNP GLs only, or both SNP and SV GLs, respectively.

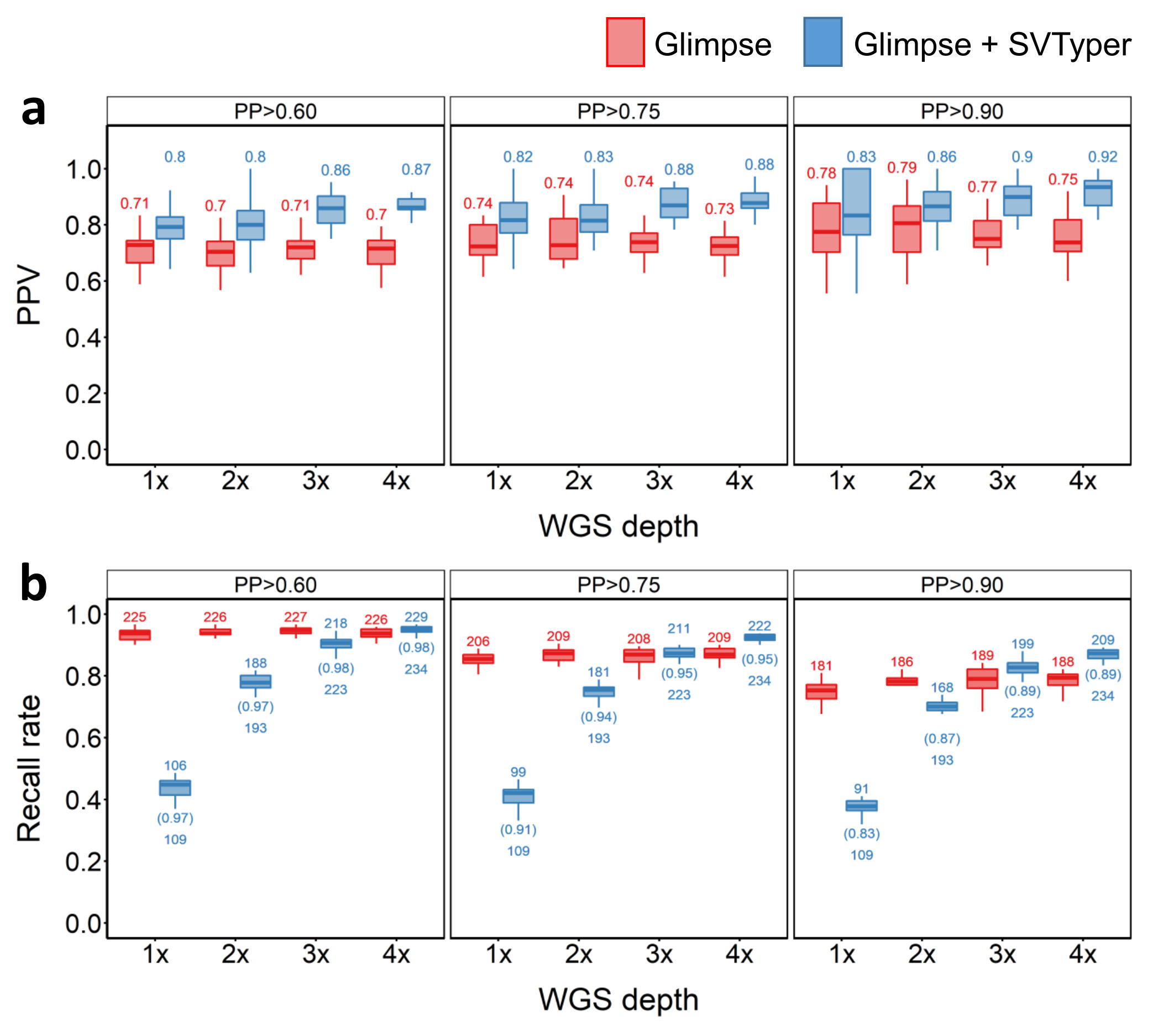

**Supplementary Figure 16. Performance of out-panel sample (n=20) inversion SV imputation.** PPV (**a**) and recall rate (**b**) for across different PP cut-offs, sequencing depths and strategies used. Red and blue plots indicate PPVs when the pipeline was supplied with SNP GLs only, or both SNP and SV GLs, respectively.
